# Assembling unmapped reads reveals hidden variation in South Asian genomes

**DOI:** 10.1101/2025.05.14.653340

**Authors:** Arun Das, Arjun Biddanda, Rajiv C. McCoy, Michael C. Schatz

## Abstract

Conventional genome mapping-based approaches systematically overlook genetic variation, particularly in regions that substantially differ from the reference. To explore this hidden variation, we examined unmapped and poorly mapped reads from the genomes of 640 human individuals from South Asian (SAS) populations in the 1000 Genomes Project and the Simons Genome Diversity Project. We assembled tens of megabases of non-redundant sequence in tens of thousands of large contigs, a significant portion of which is present in both SAS and non-SAS populations. We demonstrated that much of this sequence is not discovered by traditional variant discovery approaches even when using complete genomes and pangenomes. Across 20,000 placed contigs, we found 8,215 intersections with 106 protein coding genes and >15,000 placements within 1 kbp of a known GWAS hit. We used long read data from a subset of samples to validate the majority of their assembled sequences, aligned RNA-seq data to identify hundreds of unplaced contigs with transcriptional potential, and queried existing nucleotide databases to infer the origins of the remaining unplaced sequences. Our results highlight the limitations of even the most complete reference genomes and provide a model for understanding the distribution of hidden variation in any human population.

## Introduction

Genome sequencing studies have not uniformly spanned the spectrum of human diversity. As of 2019, >80% of participants in genomic studies were of predominantly European ancestries, with poor representation from other geographic regions ^1,2^. While recent sequencing and assembly efforts are now addressing this gap ^3–5^, the persisting biases hinder fundamental biological research and clinical interpretation ^6,7^.

One such underrepresented region is South Asia, home to more than 1.9 billion people ^8^, with another 35-40 million people in its diaspora ^9–11^. Population genomic research focused on South Asia has uncovered deep population structure, runs of homozygosity ^12^ attributed to founder effects and endogamy, as well as intricate histories of both ancient and recent admixture, all of which are influenced by societal and cultural factors ^13–17^. Although these studies are invaluable, South Asians compose only ∼2% of individuals in genome-wide association studies (GWAS), and even within this broader group, certain ethnicities are over- or underrepresented ^1^ (**Supplementary Figure 1**). Prior efforts to catalog South Asian diversity are largely out-dated, based on genotyping arrays or low-coverage sequencing data, or performed using historical versions of the reference genome^18,19^. While there is an abundance of low-coverage sequencing data from individuals of South Asian ancestries, curated, high-coverage, open-access datasets from South Asian samples are lacking. Two notable exceptions are the 1000 Genomes Project (1KGP) and Simons Genome Diversity Project (SGDP), which offer high-coverage whole-genome sequencing data from 640 South Asian individuals across 24 populations (Figure 1a-b).

**Figure 1.**
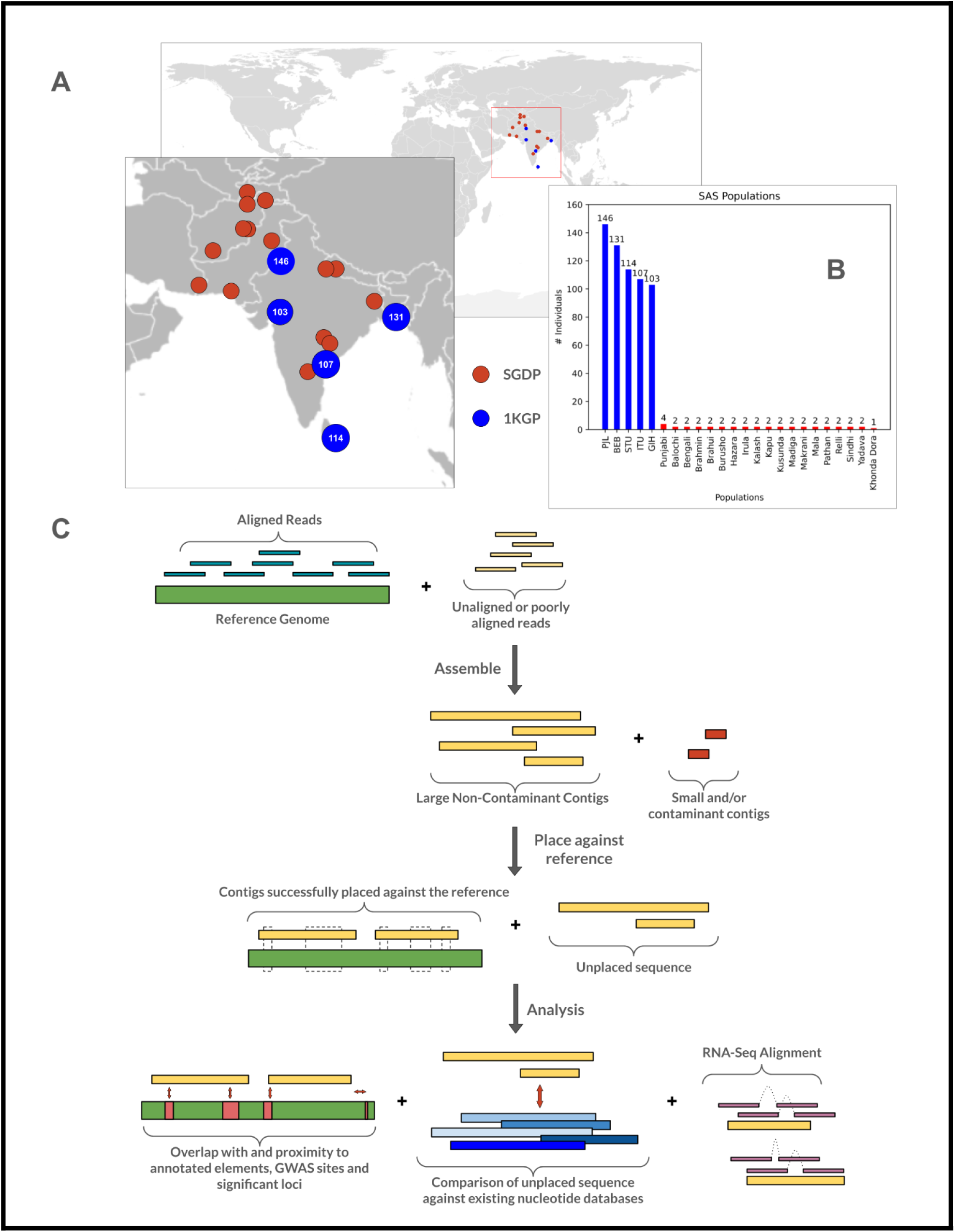
Input Data and “Read Rescuing” Pipeline. **(a)** Geographic locations of populations from which data were sourced; red dots indicate samples from SGDP, while blue dots indicate samples from 1KGP. **(b)** Number of individuals in each of the 24 SAS populations. **(c)** Schematic of the analysis pipeline. Existing short read data were aligned against a reference, after which the unaligned reads were extracted and assembled into larger contigs, in turn realigned to a reference genome. Successfully placed contigs were tested for overlaps and proximity with annotated genomic elements (e.g., protein-coding genes) and GWAS hits, while unplaced loci were queried against existing nucleotide databases. A full pipeline overview is available in **Supplementary Figure 2**.

Even with high-quality data, researchers face additional biases that impede genetic studies. Most conventional genomic analyses involve alignment to a reference genome, but a single reference genome may obscure variation in regions poorly represented by the reference. Even for the complete T2T-CHM13 reference genome, insertions relative to the reference and highly divergent haplotypes may fail to align ^4,5,20^. This reference bias was highlighted in the African Pangenome project (APG) ^21^, which found hundreds of megabases of sequence present in African individuals that were absent from GRCh38. Similar efforts with other populations have similarly identified tens of megabases of sequence that are missing in widely used reference genomes ^22–24^. Even current pangenomes cannot fully overcome this challenge, as human populations possess an excess of ultra-rare variation that will not be adequately captured by any modest-sized panel of references ^3,25^.

In this work, we cataloged genetic diversity within South Asian populations and tested the limitations of reference genomes and pangenomes. To this end, we re-analyzed existing short read sequencing data from 640 South Asian individuals across 24 populations in the 1KGP ^26,27^ and SGDP ^28^ datasets and assembled sequences that are present in these samples but absent from reference genomes **(Figure 1c)**. This process involved assembling unmapped reads into larger contigs, which has been demonstrated to reveal novel and variant sequences in both human and non-human samples ^21,29–33^. We then validated thousands of large contigs assembled from these unmapped reads through alignment to long-read based assemblies ^34^, effectively anchoring approximately 20,000 contigs onT2T-CHM13 and facilitating interpretation. Many of these placements overlapped with biologically relevant regions, including thousands of intersections with protein coding genes or in proximity to GWAS hits. In contrast, conventional pipelines failed to detect and place most of these large contigs and ignored most of the unplaced novel sequence. Through alignment of existing RNA-seq data, we also uncovered several contigs with transcriptional evidence, underscoring their functional and phenotypic potential. Together, our research highlights the limitations of both single and pangenome references and presents a generalizable framework and open-source software pipelines for recovering hidden variation from existing sequencing data.

## Results

### Alignment, assembly, and filtering

#### Alignment and assembly of data from the 1KGP SAS set

We first aligned reads from the 601 1KGP SAS individuals against the T2T-CHM13 v2.0 ^4^ and GRCh38 ^35^ linear references. Alignment to the newer T2T-CHM13 reference improved the alignment rate (+0.7-1.0%) compared to GRCh38 and increased the number of reads concordantly aligned to a single locus (+1.0%) **(Figure 2a)**. This improvement was expected, since T2T-CHM13 adds hundreds of millions of bases of newly resolved sequence. However, despite this improvement, 1.3-1.7% of all reads per individual remain unaligned against T2T-CHM13. These reads represent an opportunity for discovery, as they may contain novel sequences these individuals possess relative to the chosen reference, as well as reads that fail to align due to sequencing errors or other technical artifacts. In addition to extracting fully unaligned reads, we also examined reads with poor alignment quality scores (MAPQ < 20), which constituted 20-30x the number of fully unaligned reads. Results from the alignment and assembly of these reads are provided in the **Supplementary Materials**.

**Figure 2.**
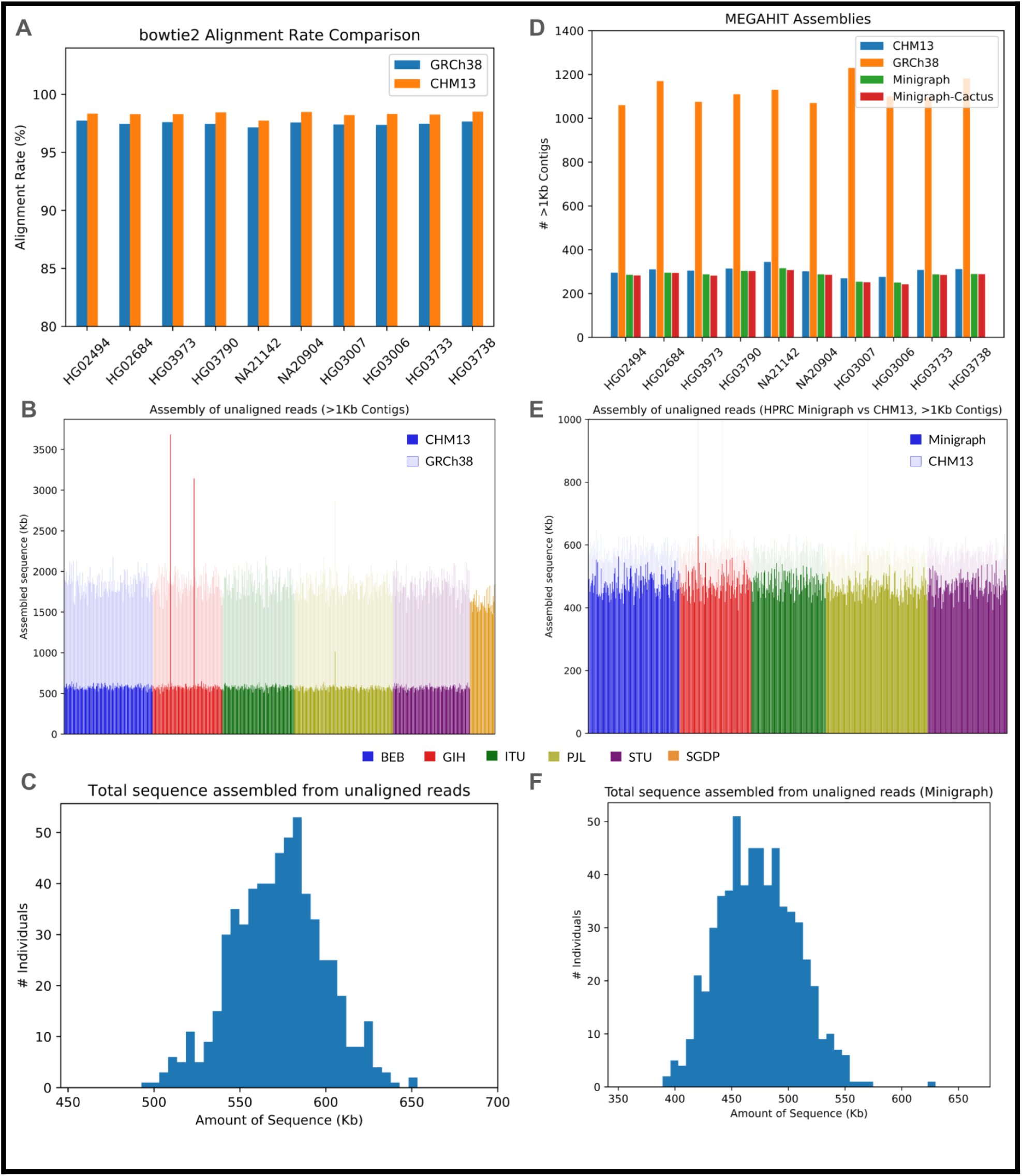
Genome Alignment and Assembly. **(a)** Comparison of the bowtie2 alignment rate between GRCh38 and T2T-CHM13 across 10 SAS samples. **(b)** Amount of assembled sequence from unaligned reads against linear references per individual across the entire 1KGP and SGDP SAS sets. For the 1KGP subset, results are depicted against both T2T-CHM13 (solid) and GRCh38 (translucent) reference genomes. **(c)** Amount of assembled sequence from unaligned reads against T2T-CHM13 across the 1KGP SAS individuals. **(d)** Comparison of the number of assembled >1 kbp contigs across T2T-CHM13, GRCh38, and the HPRC constructions across the same 10 SAS samples. **(e)** Amount of assembled sequence from unaligned reads against the Minigraph pangenome reference per individual across the entire 1KGP SAS set. The T2T-CHM13 results are depicted with translucent bars and displayed behind the solid bars for comparison. **(f)** Amount of assembled sequence from unaligned reads against the Minigraph pangenome reference across the 1KGP SAS individuals.

Unaligned reads were then assembled into larger contigs whose origins could be more easily assessed. Using the MEGAHIT genome assembler ^36^ on reads that failed to align against T2T-CHM13, we initially assembled ∼1 Mbp of sequence per individual. However, much of this sequence was composed of short contigs, potentially reflecting contamination or assembly errors. Using an approach inspired by the African Pangenome study ^21^, this set was filtered to exclude contigs <1 kbp in length. This reduced the amount of assembled sequence to approximately 550 kbp per individual. These contigs were then screened using BLAST ^37^ and Centrifuge ^38^ against databases of known bacterial and microbial sequences, removing contigs with any matches to these contaminants. This step removed few contigs (< 0.5%), but increased confidence that the remaining sequences were of human origin. Of the contaminant contigs that were removed, the majority matched to adapter sequences or common human contaminants such as herpes virus.

After filtering, we assembled an average of ∼550 kbp of sequence per SAS individual, totaling approximately 350 Mbp across the entire 1KGP SAS set (**Figure 2b-c**). While three outlier individuals were apparent, self-alignments of their contigs revealed nearly-duplicated assembled sequence, thereby resolving these cases (**Figure 2b**). Across all samples the size of these contigs varied greatly, with an average of 70 contigs >5 kbp and 18 contigs >10 kbp per individual. The average N50 for these filtered assemblies was 1.95 kbp, and the average size of the largest contig assembled per individual was 15-18 kbp, with the largest contigs in the entire set spanning over 50 kbp. The analogous approach using GRCh38 assembled an average of 2 Mbp of sequence per individual, significantly exceeding the above results from T2T-CHM13 (two-tailed Welch’s t-test, t = -134.45, p-value < 1 × 10^−308^), and underscoring the greater completeness of the T2T reference.

Application of RepeatMasker ^39^ showed that 4.39% of the assembled sequence consisted of short interspersed nuclear elements (SINEs) (1.73% ALUs, 2.66% MIRs), 17.89% long interspersed nuclear elements (LINEs) (13.85% LINE1, 3.89% LINE2, 0.11% L3/CR1), 8.54% long tandem repeat elements (LTRs) (3.40 ERV-L, 3.52% ERVL-MaLRs, 1.04% ERV Class I, 0.04 ERV Class II) and 2.53% DNA elements, together constituting a total of 33.40% interspersed repeats (**Supplementary Table 8**). An additional 2.2% of sequence consisted of small RNA (0.03%), satellites (0.64%), simple repeats (1.21%), and low complexity DNA (0.32%). In comparison, the contigs assembled from GRCh38 unmapped reads contained substantially more SINEs (8.5%), satellites (35.5%) and simple repeats (17%), but a lower proportional composition (but greater absolute composition) of LINEs (7.5%), LTR elements (3.56%) and DNA elements (0.9%).

We further evaluated the impact of the complete T2T-CHM13 reference by performing “cross alignment”, where the contigs assembled from T2T-CHM13 unmapped reads were aligned against those assembled from GRCh38 unmapped reads (**Supplementary Figure 3**). We observed that the T2T-CHM13 contigs were largely a subset of the GRCh38 contigs, with 70% having a high identity, high mapping quality alignment against a GRCh38 contig. Conversely, 75% GRCh38 contigs did not align well to the T2T-CHM13 set, but 70% aligned well to the T2T-CHM13 reference itself, reflecting its completeness and mirroring results from the African Pangenome study ^21^. Our results suggest that use of GRCh38 obscures genetic variation, particularly within erroneously assembled or unresolved regions that were later resolved in T2T-CHM13. For these reasons, unless otherwise noted, we focused our subsequent analyses on the contigs assembled from T2T-CHM13-unaligned reads.

#### Alignment and assembly of data from the SGDP SAS set

We repeated the alignment and assembly steps with data from 39 SAS individuals across 19 populations from the SGDP dataset. For 37 out of the 39 individuals, assembly of their unaligned reads against T2T-CHM13 (0.9 - 1.7% of the total input reads) and filtering to >1 kbp non-contaminant contigs resulted in 1.4 - 1.8 Mbp of assembled sequence per individual. Each assembly contained ∼500 contigs with average N50s of 3.2 kbp, and the largest assembled sequences reached ∼40 kbp (**Supplementary Figure 4**). We attribute the larger span of non-contaminate contigs to the higher sequencing depth of SGDP (32-66x, average coverage of 43x) compared to 1KGP (∼30x).

Two outlier individuals from the Kusunda group exhibited markedly different alignment and assembly results. Specifically, ∼10% fewer reads from these samples aligned to the reference and the unaligned reads assembled into >200x as much sequence as other SGDP samples (**Supplementary Figure 5)**. BLAST and Centrifuge did not categorize these sequences as known human sequence or contaminants, and most quality metrics were in line with other SGDP SAS samples. Full details of these two outliers are provided in **Supplementary Figure 6**, but we hypothesize that the excess sequences originate from a mixture of unknown microbial contaminants and sequencing artifacts, and we therefore excluded these two samples from subsequent analyses.

#### Pangenomic improvements in alignment and assembly

We repeated the assembly process with reads that failed to align to the HPRC Minigraph and Minigraph-Cactus v1.0 pangenome references ^3^. These HPRC pangenome references were constructed by augmenting the T2T-CHM13 reference with the genome sequences from 47 individuals, including one individual from the South Asian continental group ^3^. Against the Minigraph reference, alignment rates rose by 0.3-1.1% per individual, which in some cases halved the number of unaligned reads compared to results using T2T-CHM13. The expanded Minigraph-Cactus reference further improved the alignment rates, though this improvement was marginal (<0.1%). For the SAS individual also present in the Minigraph and Minigraph-Cactus reference, we observed near perfect alignment across the genome, with the overall alignment rate rising from 98.7% against T2T-CHM13 to 99.8% against the Minigraph reference.

Despite the sizable declines in the number of unaligned reads, the reductions in the amounts of assembled sequence were not as dramatic (**Figure 2d-f**). Using reads that failed to align to the Minigraph pangenome reference, we assembled 450-500 kbp of sequence per 1KGP SAS individual in >1 kbp non-contaminant contigs, a ∼10% decrease in sequence compared to the same individuals using T2T-CHM13 (Welch’s t-test, two-tailed p-value of 2.03 × 10^−13^; **Figure 2e-f**). This proportionally modest decline suggests that many of the newly aligned reads would have assembled into short contigs discarded during our filtering step. The number of >1 kbp non-contaminant contigs declined even less (**Figure 2d**), suggesting parts of previously assembled contigs had been resolved. The average N50 of the filtered assemblies was 1.9 kbp and the largest assembled contigs per individual were 15-18 kbp (maximum ∼50 kbp). The augmented Minigraph-Cactus reference resulted in further modest declines in the amount of assembled sequence (<5%) and number of contigs that composed this sequence (<3.5%).

#### Comparison to non-SAS populations

To contextualize these results, we compared to analogous results obtained from samples from non-SAS populations. Specifically, we repeated the same alignment and assembly steps using 10 randomly selected individuals, 5 XX and 5 XY, from each of the 21 non-SAS populations in the 1KGP dataset. Across the two linear references, the results with the non-SAS set were similar to those from the SAS set, with approximately three times as much sequence assembled per individual from reads that failed to align to GRCh38 versus reads that failed to align to T2T-CHM13. However, individuals from the SAS set produced on average 10-15% less assembled sequence than individuals from the seven African (AFR) populations (two-tailed Welch’s t-test; t = -36.8; two-tailed p-value = 1.103 × 10^−46^), and on average 5% more sequence than the individuals from the non-AFR, non-SAS populations ((two-tailed Welch’s t-test; t = 6.22; two-tailed p-value = 5.39 × 10^−9^) (**Supplementary Figure 7, Supplementary Table 9**). Performing population level comparisons, where the sequence assembled across 601 SAS individuals are compared just to the 10 individuals from each non-SAS population, we see all seven AFR populations have a two-tailed p-value < 1.76 × 10^−7^, and the non-SAS populations CHS, CDX, PEL, GBR, FIN and CEU have a two-tailed p-value < 0.05 (**Supplementary Table 9**).

It is worth noting that while T2T-CHM13 exhibits the greatest genetic similarity to 1KGP samples from the European (EUR) continental group, the amount of sequence assembled for samples from EUR populations is nearly the same as any other non-AFR population. This underscores the fact that human genetic diversity tends to be either globally common (present in all populations at relatively similar frequencies) or rare (geographically restricted, but only because it is rare and is also rare within a given population) ^40^. More broadly, these results are consistent with previous knowledge that 1) genetic diversity in African populations exceeds that in non-African populations as a result of the out-of-Africa bottleneck ^41,42^ and 2) there is an abundance of rare variation in human genomes due to rapid population growth ^43^.

Overall, these results highlight the improvements offered by pangenomes, which recover common variation that previously failed to map. However, substantial amounts of sequence can be assembled from reads that fail to align even to these pangenome references, highlighting the need for alternative approaches beyond reference-based alignment.

### Placement of assembled contigs

Placing assembled contigs back into the reference genome may be possible through direct alignment of the contigs to the reference genome or using data from mapped mate pair reads ^21^. Using the direct alignment approach, approximately 40% of contigs aligned at least partially to the T2T-CHM13 reference, declining to ∼8% upon required alignment lengths of at least 500 bp. This equated to an average of ∼350 contigs that remained unaligned per individual, and we hypothesized that these may contain any entirely novel sequence. In contrast, the mate pair approach placed fewer contigs than direct alignment (3,952 / 1.9%), but addressed cases where contigs diverged from the reference sequence. Combining these two methods, 19,691 of 199,564 contigs (9.8%) >1 kbp in length were placed against T2T-CHM13, with enrichments noted on chromosomes 6, 9, and Y (**Supplementary Figure 8**), potentially driven by their higher repeat content (e.g., the HLA locus on chr6). Of the placed contigs, and restricting to the 21 SAS individuals with available long-read data, >85% could be aligned with high identity against a personalized long read assemblies. While extending across all chromosomes, these placements were not evenly distributed throughout the genome, with enrichments noted on chromosomes 6, 9, and Y (**Supplementary Figure 8**), potentially driven by their higher repeat content (e.g., the HLA locus on chr6).

### Intersection with annotated elements

We intersected the placed contigs with the “JHU RefSeqv110 + Liftoff v5.1” gene annotations from the T2T Consortium using bedtools ^44^. Across the SAS set, we identified 52,727 intersections with known transcripts, 12,021 with known genes, 5,050 with known exons, and 3,015 with known CDS regions (**Figure 3a**), with substantial heterogeneity noted across the genome (**Figure 3b**; **Supplementary Figure 9**). These intersections affected a total of 1,189 unique transcripts, 172 genes (106 protein-coding), 33 exons and 6 CDS regions (**Figure 3c**). While the majority of the 106 protein coding genes intersected with contigs from one or few individuals, some reflected more common variation (**Supplementary Figure 10**). Qualitatively similar results were observed when applying the same approach to samples from non-SAS populations (**Figure 3d**). Gene intersections in the SAS set included 11 genes where mutations have been related to eye disorders (*AGBL1, CPAMD8, CRB1, HMCN1, KIF14, LIN9, MYO7A, PPEF1, TTC39C, USH2A* and *WDR17*), as well as genes involved thyroid function (*TSHR*), facial dysmorphia (*GPC3, CPA6, OPHN1, OR51A4* and *SNP1*), ciliary function (*CC2D2A, DRC1, NME8, TTC39C*), and cancers (*DLEC1, DMBT1, LIN9, PTPRO, SEM1, SYK and ZNF292* **Supplementary Table 1**; **Supplementary Table 6**-**7**). The observed count of 12,021 gene intersections was substantially lower than the number of intersections obtained through random placement of contigs against the reference genome (14,500 intersections, averaged over three runs), consistent with the action of purifying selection against common structural variation in genic versus non-genic regions.

**Figure 3.**
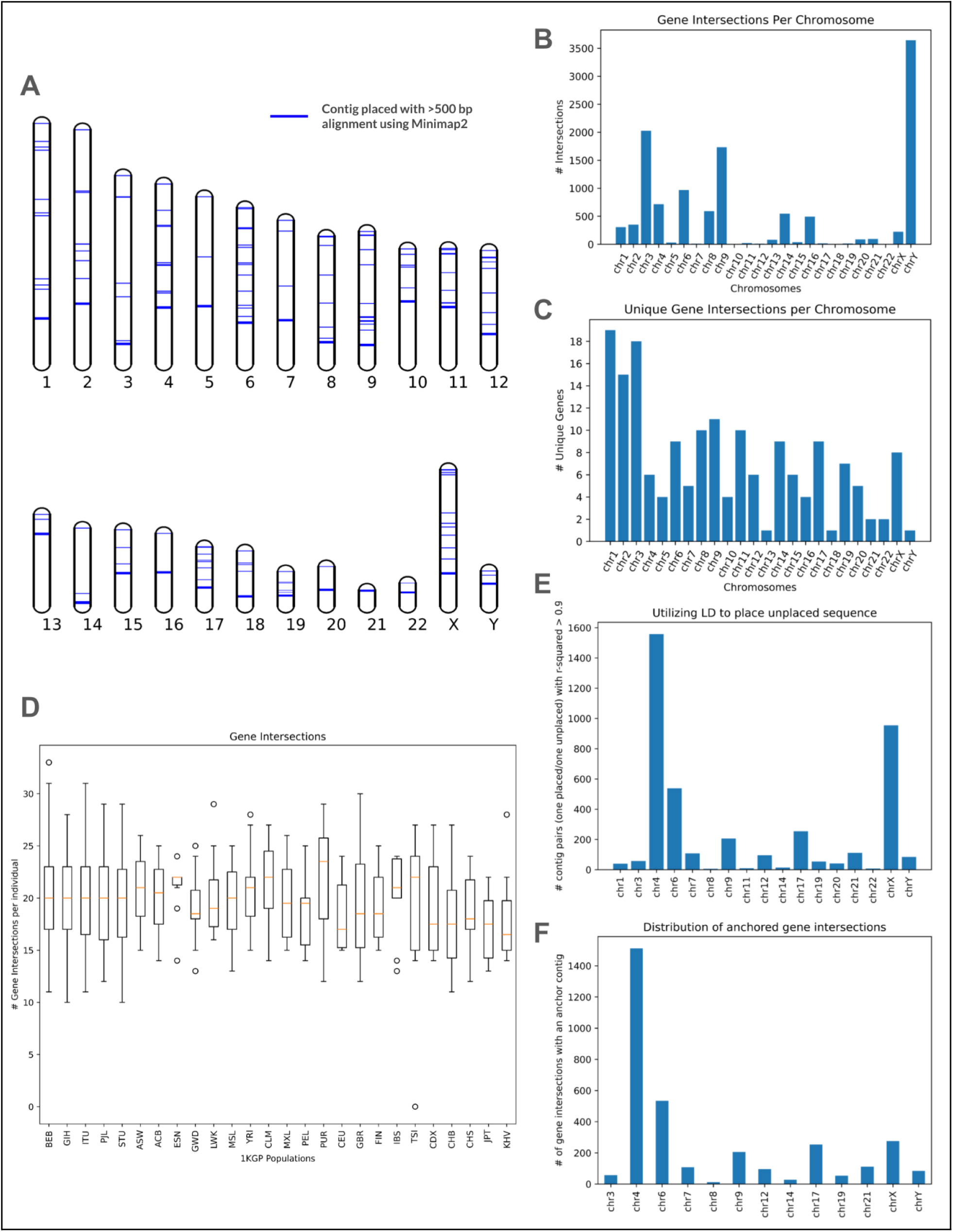
Placements & regions with “hidden” SA variation. **(a)** Visualization of placements against the T2T-CHM13 reference. Each line represents an assembled contig with a >500 bp Minimap2 alignment against the T2T-CHM13 reference. **(b)** Counts of the number of gene elements in T2T-CHM13 v2.0 intersecting placed contigs. **(d)** Distribution of the number of gene intersections per individual across the 26 1KGP populations. **(e)** Distribution of contigs anchored via LD (R^2^ > 0.9) with a placed contig. **(f)** Distribution of gene intersections with anchored contigs, where a pair of contigs have R^2^ score > 0.9, and one of the pair is placed and the other is unplaced.

#### Linkage disequilibrium within the shared contig set

The presence/absence of each shared contig across all 640 SAS individuals allowed us to measure linkage disequilibrium (LD) between pairs of contigs and identify “blocks” of placed contigs that likely belong to the same larger sequence. Specifically, we computed the squared Pearson correlation coefficient (r^2^) between all pairs of 13,875 shared contigs and found 644,578 pairs of contigs with r^2^ > 0.9 (0.33% of all pairs). Of these, 194,428 (0.1% of all pairs, 30% of pairs with r^2^ > 0.9) are between contigs on the same chromosome and 56,935 are between contigs within 10 kbp of each other. We also observed 4,142 pairs where a placed contig is in strong LD with an unplaced contig (r^2^ > 0.9 across the 640 individuals). In these cases, the placed contig can act as an “anchor”, allowing us to narrow down the potential location of the unplaced contig within the genome. Within these anchor contigs, we identified 3,334 intersections with 24 genes across 13 chromosomes (**Figure 3e-f**), as well as 58 intersections with known GWAS hits.

#### Analysis of unplaced sequence

Despite placing approximately 20,000 contigs from the SAS set against T2T-CHM13, we assembled over 180,000 contigs that could not be placed. By querying this set of unplaced contigs using BLAST, we found that >90% had a high similarity match with non-reference human sequence or reference and non-reference primate sequences (**Supplementary Table 2)**, supporting their non-contaminant human origin. We also observed a small fraction of contigs with high similarity to non-primate mammalian sequences, potentially reflecting highly conserved loci.

#### Analyzing pangenome placements

The limited availability of pangenome tools precluded extension of the placement approach to the graph pangenomes. As an alternative, we performed conventional alignment to the linear references that comprise the pangenome, identifying 8.5% of the assembled contigs (16,854/199,564) with an alignment length >500 bp, thus slightly exceeding the number of contig alignments to T2T-CHM13 alone (15,739).

Of the contigs that failed to be placed against T2T-CHM13, 905 of 179,873 (0.5%) aligned to the HPRC Minigraph pangenome. These alignments occur throughout the genome, with no specific chromosome or region especially enriched for these placements. These results highlight the power of pangenome approaches while emphasizing the need for centralized annotations and auxiliary data to facilitate downstream interpretation.

### Validation with long reads

The initial data release from the 1KGP ONT effort <u>(Gustafson et al. 2024)</u> contains long read data from 21 individuals present in our SAS set, which can be used to help validate the assembled contigs from these individuals. We constructed simple assemblies for these 21 individuals with their long reads using the Flye assembler (Kolmogorov et al. 2019). The assemblies contain 2.81 to 2.84 Gb of sequence, with N50s ranging from 31.9 to 49.2 Mb and mean coverage of 29 to 47x. Although these assemblies are far from telomere-to-telomere completeness, we observed that 40-50% of these previously unaligned reads aligned well to the new personalized long read assemblies of the respective samples. The “rescue” of these unaligned reads highlights the limitations of a single reference genome and confirms that much of the missing sequence is variation from loci that are easily assembled from long-reads.

For the same 21 samples, we also aligned the contigs we previously assembled from reads that failed to align to T2T-CHM13 to the corresponding long-read assemblies, finding that 85% of contigs aligned with high identity (**Figure 4a**). These included 92.7% (331/357) placed contigs that intersect with genes (**Figure 4b**). We hypothesize that the remaining 15% of contigs originate from more complex regions that may require tailored assembly approaches or additional sequencing with complementary platforms.

**Figure 4.**
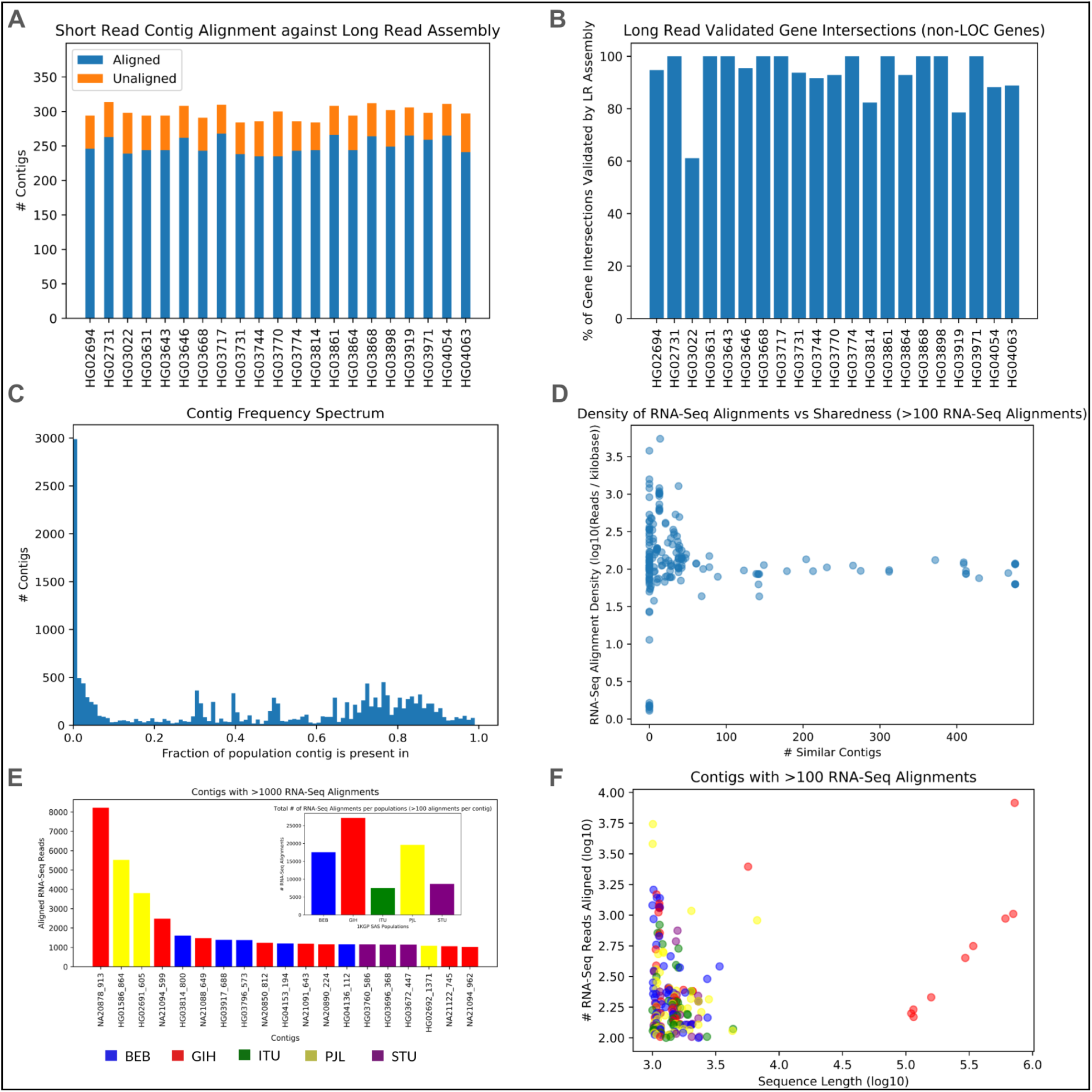
RNA-Seq Alignment, Long Read Validation and characterizing shared sequence. **(a)** The number of unaligned read contigs in 21 SAS individuals that are successfully aligned against a personalized long-read reference from the same individual**. (b)** The fraction of gene intersections from each of 21 SAS individuals that are validated based on alignment to long-read assembled genomes. **(c)** Allele frequency distribution of contigs, determined through all-by-all alignment and assessment of sequence similarity. **(d)** How common the most aligned-to contigs from RNA-Seq analysis are, plotted against the density of RNA-seq alignments (number of reads aligned per kilobase of contig, log_10_). **(e)** The 20 contigs across the 140 SAS individuals with the most aligned RNA-Seq reads , colored according to their SAS subpopulation. These contigs come from 19 individuals across the SAS set. Inset in (a) is the number of contigs with >100 RNA-Seq alignments in each of the five SAS sub-populations. **(f)** All contigs with >100 RNA-Seq alignments during all-vs-all alignment, with the number of alignments plotted against their sequence length, colored according to their SAS subpopulation.

We repeat this long read validation using a second dataset from Schloissnig et al. (Schloissnig <u>et al. 2025)</u>, and perform the same analysis on 20 SAS individuals from this set. The Flye assemblies for these individuals are comparable to those from the 1KGP set (2.65 to 2.82 Gb of sequence, N50s of 25.6 to 45.7 Mb, and mean coverage of 14 to 33x), and initial alignment sees 40-50% of previously unaligned reads now align to the personalized long read assembly. Contig alignment is slightly lower with this data (80-87%), but in line with the fact this dataset is substantially lower coverage than the 1KGP ONT set.

### Shared sequence analysis

Summing over all SAS samples, we assembled 410 Mbp of sequence in > 1 kbp, non-contaminant contigs from short reads that failed to align to T2T-CHM13. However, we anticipated that much of this sequence would be redundant, as multiple individuals may share an allele (identical by descent, or more rarely, by recurrent mutation) that is divergent from the reference (**Supplementary Figure 11**). To address such cases, we performed an all-vs-all alignment of the assembled contigs using Minimap2 ^45^, and considered contigs with >90% alignment to have originated from a shared allele. In total, this procedure collapsed the 199,564 contigs totalling 410 Mbp of sequence into 13,875 shared contigs and 2,137 unique (i.e., “non-redundant”) contigs totalling 47 Mbp of sequence. The allele frequency spectrum demonstrates that a large fraction of the contigs are rare (present in <10% of the combined SAS set), with the majority of these present in just 1 or 2 individuals, consistent with the known abundance of rare variation in human populations. (**Figure 4c**, **Supplementary Figure 12-13**). We note that the presence of trios distorts the frequency spectrum compared to expectations for an unrelated sample, but the practical effect is minimal in practice (e.g., enriching for doubletons versus singletons).

We were also interested to quantify the amount of sequence from these SAS samples that is present in genomes of individuals from non-SAS populations. To quantify this proportion, we aligned read data from the selected 210 non-SAS individuals against the non-redundant SAS contigs. We found that, on average, 3.8 - 4.5% of reads from each individual align to this set of contigs, with individuals of all 21 non-SAS populations exhibiting roughly the same alignment rate. This observation further highlights the fact that most variation is rare, and that the variation that is common tends to be shared across all populations.

### Comparison to traditional variant finding approaches

To evaluate the strengths and limitations of our “rescued read assembly” approach, we compared our results to those obtained from conventional short-read-based SV callers, starting with Manta ^46^ and LUMPY ^47^ and focusing on insertions (given that these are also the focus of our assembly strategy).

Manta identified an average of 2,300 insertions per individual, but only an average of nine insertions per individual were longer than 500 bp (192 in total, with a maximum of 16 found in HG03898), and no insertions were longer than 1 kbp. In contrast, our approach assembled an average of 299 (SD = 9.3) contigs longer than 1 kbp per individual, an average of 36 of which could be placed against T2T-CHM13. The placed contigs overlapped some of the smaller and many of the larger (>500 bp) insertions called by Manta but also included hundreds of placed sequences that were not detected by Manta (**Supplementary Figure 14-15**), 85% of which were validated by long reads. LUMPY is largely focused on deletion calling and is limited in its ability to detect small insertions, but is able to detect large insertions through identification of breakends ^47^. Across the 21 SAS individuals, LUMPY detected none of these large insertions, despite our approach identifying and placing several large contigs in these samples, further validated with long reads.

We also compared the sequence assembled by our approach to the sequences found by PopIns2 ^48^, which detects large non-reference sequences by assembling reads that do not have a high quality reference alignment. Across a random selection of 10 1KGP individuals (one XX and one XY from each of the five SAS 1KGP populations), PopIns2 assembled an average of 6,264 contigs >1 kbp per individual from reads that failed to align to T2T-CHM13, in broad accordance with our results (**Supplementary Results, Supplementary Figure 16**). Comparing the placement of these contigs against the contigs assembled and placed by our approach, we found that of the 267 contigs placed against these 10 individuals, 210 overlapped a placed Popins2 contig. However, our approach assembled more large contigs than PopIns2 (average of 510 contigs >5 kbp and 95 contigs > 10 kbp by our approach vs 432 contigs >5 kbp and 72 contigs >10 kbp by PopIns2) and also places a larger fraction of such contigs (10-12% of our contigs vs 8% of PopIns2 contigs) based on alignment, LD, and mapped mate pair reads.

### Functional analysis of hidden variation

#### RNA-seq analysis

We used RNA-seq data from the MAGE project ^49^, obtained from lymphoblastoid cell lines from 731 globally diverse individuals from 1KGP, to evaluate the functional potential of the previously assembled contigs. We specifically focused on the 140 MAGE samples from SAS populations that overlap with our study. We aligned RNA-seq reads from each individual to their contigs that were assembled from reads that failed to align to T2T-CHM13. Per individual, ∼800-1,000 reads aligned to their respective contigs, with the majority (>80%) spread across few contigs. While this represents a small fraction of the input RNA-seq dataset (<0.1%), this is not unexpected, as the total length of assembled contigs per individual is <0.02% the size of the human genome. Across the 140 individuals, 90% of contigs with >100 RNA-seq alignments had BLAST hits to non-reference human and primate sequences, some of which is annotated as functional (**Supplementary Table 3-4**). These include hits to annotated non-reference sequences from chromosomes 1, 2, 3, 7, 9, 15, X and Y.

We also aligned the RNA-Seq data from all 140 individuals against a combined set of filtered contigs from all 601 1KGP SAS individuals and identified 200 contigs with >100 read alignments and 19 contigs with >1,000 read alignments (**Figure 4e-f, Supplementary Figure 17-19**), 83% of which had BLAST hits to non-reference human and primate sequences (**Supplementary Tables 3-4).** Across both individual-specific and combined RNA-seq alignments, we found that the majority (>80%) of highly aligned-to contigs appear in just one or a few individuals, with few classified as widely shared sequences (**Figure 4d**).

Intersecting the placement information for these contigs with the RNA-seq alignments, we find that the majority of highly aligned-to contigs could be placed via alignment to T2T-CHM13 (140/200 of contigs with >100 alignments, 12/19 contigs with >10 alignments), with most of the placements tracing to the Y chromosome. Aligning these contigs against the HPRC Minigraph pangenome reference modestly increased the rate of alignment (144/200 of contigs with >100 alignments, 13/19 contigs with >10 alignments), with few additional contig alignments to chromosomes 19 and Y.

#### Intersection with GWAS hits

We next explored the phenotypic relevance of the assembled contigs by comparing the placed sequences against existing catalogs of trait-associated loci. We intersected the set of placed contigs against the GWAS v1.0 catalog (lifted over to T2T-CHM13 coordinates ^4,5^) and identified 2,812 cases where a placed contig overlapped an annotated GWAS hit. These hits were largely concentrated on chromosome 6 (1,995), Y (601), and 5 (208), with small counts noted on other autosomes (**Figure 5b**). Loosening our search criteria, we identified >6,000 contigs within 100 bp of 36 unique GWAS hits and >15,000 contigs within 1 kbp of 112 unique GWAS hits (**Figure 5a-b**), the vast majority of which were validated with long-read data (<100 bp distance: 215/254, 84.6%; <1 kbp distance: 506/587, 86.2%; **Supplementary Figure 20**).

**Figure 5.**
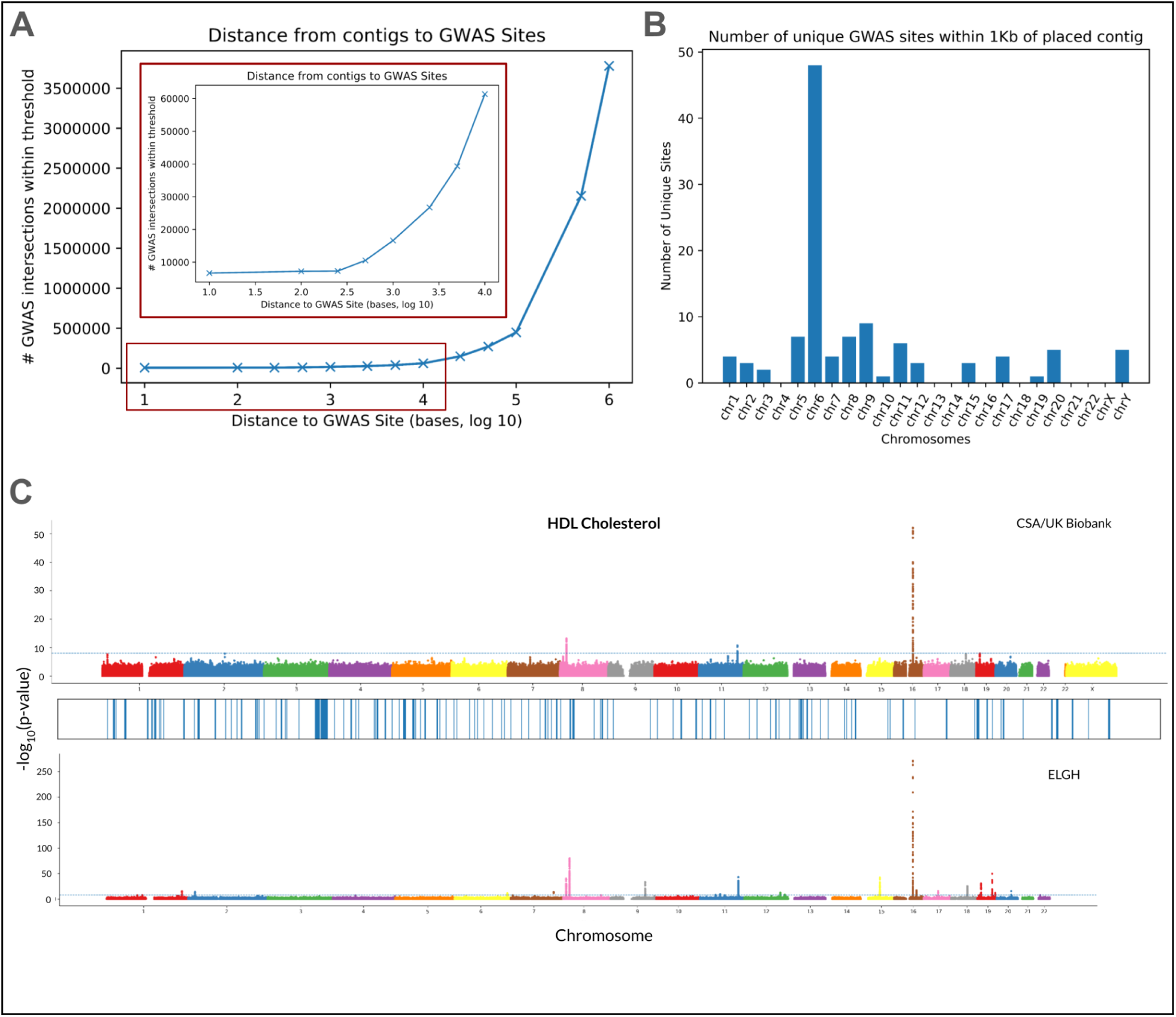
Functional Analysis. **(a)** The distance between a placed contig and a known GWAS site across a range of thresholds. **(b)** The number of unique GWAS sites with a placed contig within 1 kbp across the 24 chromosomes. **(c)** Significant loci (p-value < 1✕ 10^−8^) associated with HDL cholesterol in the CSA population of the UK biobank (**top**) and the entire ELGH cohort (**bottom**), alongside locations of placed sequence against GRCh38 (**middle**).

Finally, we investigated if any of the placed contigs overlapped with or were near trait associated loci from the UK Biobank (UKB) ^50,51^ and East London Genes & Health (ELGH) study ^52^ (**Figure 5c**). The former contains data from ∼400,000 individuals of all ancestries, while the latter contains data from ∼37,000 SAS individuals in the UK. Across the 21 biomarker traits from the Pan-UK Biobank ^53^ and 42 biomarker traits from the the Genes and Health (ELGH) ^52^ dataset, we identified only 5 placed contigs within 10 kbp of significant locus at a relaxed p-value threshold of 1 × 10^−6^. Expanding to a 1 Mbp window surrounding GWAS hits, we identified 8-14 traits close to placed contigs as we varied the p-value threshold from 1 × 10^−8^ to 1 × 10^−6^ respectively (**Supplementary Table 5**). However, we did not find any placements within 1 Mbp of a GWAS hit within the smaller UK Biobank “CSA” (Central/South Asian) ancestry group (n = ∼7,000) until we lowered our p-value threshold to 1 × 10^−5^. This is not unexpected - especially as ∼90% of our assembled contigs remain unplaced - and does not rule out the functional potential of these sequences. Understanding the functional implications of these complex loci will require future association studies based on genotype data encompassing these technically challenging loci ^54^.

## Discussion

We investigated the diversity present in genomes of South Asian human populations by aligning high-coverage short read sequencing data to reference genomes and assembling the unaligned reads. These assembled contigs contain variant sequences whose origins and functional impact we then evaluated through placement and annotation with external data sources.

We found that complete reference genomes enable better alignment, as reflected by improved alignment rates and smaller amounts of sequence assembled from unaligned reads. Despite these improvements, we still assembled hundreds of megabases of sequence (tens of megabases of non-redundant sequence), much of which cannot be placed against the reference but which we demonstrated is probably of human origin. Meanwhile, within the thousands of placed sequences, we identified thousands of overlaps with genes and variants previously associated with human traits. We used RNA-seq data to identify hundreds of contigs with functional potential, as well as long-read DNA-seq data to validate >80% of the assembled contigs and >85% of overlaps with genes and GWAS hits. We compared this assembled sequence to sequences from non-SAS populations to understand how this variation is distributed within and across populations, and we compared our results to those from existing SV calling tools, showcasing its unique advantages.

While these non-reference sequences were assembled from a South Asian sample, they do not typically exhibit strong patterns of frequency differentiation within South Asian populations. Rather, most of the variant sequences are either globally common or rare even within the South Asian populations, in accordance with the broader distribution of human genetic diversity ^55^. Together, these results highlight the abundance of genetic variation that remains undiscovered and the importance of expanding the scope of population genetic studies beyond linear reference genomes.

Further improvements in reference genomes, especially the expansion of current pangenome builds with more high quality genomes, will help to capture more diversity. The generation of more high quality DNA sequencing and genome assemblies, both within South Asia ^12,56,57^ and globally, will further expand the amount of variant sequence, while auxiliary data modalities will help validate these sequences and characterize their functional potential ^56^. Even using existing data, our analytical approach can be generalized to samples from populations around the world to discover novel genetic variation, including in groups traditionally underrepresented in genomics ^58–61^.

The variant sequences discovered using our approach can be used in multiple downstream applications. For example, the sequences could be stored as alternate sequence panels or incorporated into a tiered reference querying approach ^62^, recovering reads that fail to align against a single reference. Alternatively, the placed contigs could be used to augment pangenome builds, with the placements serving as the coordinates in the pangenome from which these sequences branch. The addition of novel sequences to the pangenome has been demonstrated to improve both variant detection and reduce variant discovery errors, motivating this application ^3^.

More broadly, our study provides a model for expanding beyond a single reference genome to more comprehensively characterize human genetic variation, establishing reference resources for the benefit of all human populations.

## Methods

### Data

We used read data from the 1000 Genomes project (1KGP) ^27^ and the Simons Genome Diversity Project (SGDP) ^28^, totaling 640 individuals from 24 South Asian populations across the two sets. The read data from these sets is high coverage (>30x), high quality, short paired reads. The 601 SAS individuals present in the 1KGP set are from five South Asian populations. Three are in the South Asian diaspora - Gujarati in Houston (GIH), Sri Lanka Tamil in the UK (STU) and Indian Telegu in the UK (ITU) - and two are in South Asia itself - Punjabi in Lahore (PJL) and Bengali in Bangladesh (BEB). The 601 individuals are largely evenly distributed between the five populations, with 109-144 individuals from each. In the 1KGP set, there are 282 XX individuals and 319 XY individuals. The SGDP set contributes 39 individuals, spread across 19 different groups within India, Pakistan, Nepal and Bangladesh. Most of these are in pairs from each group, with a single individual from one population (Khonda-Dora from India) and four from another (Punjabi from Pakistan) the only exceptions. In the SGDP set, there are 8 XX individuals and 31 XY individuals.

These reads are aligned against two linear references and two pangenome builds. The two linear references are GRCh38 ^35^ and T2T-CHM13 v2.0 ^4,20^. We used the Minigraph v1.0 and Minigraph-Cactus v1.0 builds of the newly released Human Pangenome Reference Consortium (HPRC) draft pangenome references. Both pangenome references are constructed over the T2T-CHM13 reference and include variation present in 47 selected individuals from the 1KGP set, one of which is a SAS individual present in our set. The Minigraph reference contains only SVs larger than >50bp, while the Minigraph-Cactus reference is further augmented to include base-level alignments for other variants.

We performed validation of our assembled contigs using long read data from the 1KGP ONT effort ^34^, which is available for 21 SAS individuals in our 1KGP set, and using long read data for 20 SAS individuals from the Schloissnig et al. cohort ^63^. Our functional analysis was augmented using RNA-Seq data from the MAGE dataset ^49^, which provides RNA-seq data for 140 SAS individuals. For our GWAS and Biobank analysis, we used the standard T2T-CHM13 GWAS catalog and the phenotype manifests from the Pan-UK Biobank ^53^ and the East London Genes and Health effort ^52^, focusing on the quantitative biomarker traits in each.

More details about the data characteristics of the read data and its alignments to the linear references can be found in **Supplementary** Figures 21-24.

### Read alignment and extraction

Reads from the 640 SAS individuals in the 1KGP and SGDP sets were aligned to GRCh38 and T2T-CHM13 using bowtie2 v2.4.1 ^64^, and the unaligned and poorly aligned (quality value < 20) reads were extracted using samtools v1.14 ^65^. Unaligned reads of two forms are extracted from these alignments: both reads where the read and its pair are both unaligned, as well as single unaligned reads with a mapped mate read (**Supplementary Note 1**).

### Assembling unaligned reads

We tested three assemblers on a selected group of 10 SAS individuals (two from each of the 1KGP populations): SPAdes v3.15.3 ^66^, MaSuRCA v4.0.9 ^67^ and MEGAHIT v1.2.9 ^36^. There was some variation in the number of short contigs (SPADes produces far more than the other two), but the number of large non-contaminant contigs varies by no more than 5% between assemblers. All assemblers were tested using their default parameters, with the four sets of unaligned reads extracted in the previous step passed in as input. The biggest difference between these assemblers is the variation in runtime. We opted for the fastest of these three assemblers - MEGAHIT - in an effort to speed up analysis and reduce costs (**Supplementary Note 2**).

### Filtering and processing assembled sequence

As in the APG analysis, we first discarded all contigs less than 1 kbp in size. This was done by filtering the assembly FASTA file for sequences longer than the target length, with all other sequences being discarded (**Supplementary Note 2**).

We also screened these sequences for contaminants using two tools: Centrifuge v1.0.3 ^38^ and BLAST ^37^. For Centrifuge, all assembled sequences were compared against the “Bacteria, Archaea, Viruses, Human” database, and sequences categorized as non-human are removed. With BLAST, we used the “refseq_genomic” database, and removed all sequences that have a high scoring match to a non-human sequence. The contaminant sequences can be identified by parsing the resulting outputs, and these sequences were then cut from the FASTA file. After the filtering stage, we were left with a list of >1 kbp, non-contaminant contigs from each of the SAS individuals for use in the later stages (**Supplementary Note 2**).

Repeat content in the assembled contigs and reference genomes is detected, masked and annotated using RepeatMasker v4.1.2-p1 ^39^. Using the assembled contigs as input, RepeatMasker detects and masks SINEs, LINEs, LTR and DNA elements, small RNA and satellites, and depending on the use of the “nolow” parameter, simple repeats and low complexity elements (**Supplementary Note 11**).

### Assembly of non-SAS sequences

We selected 10 individuals - 5 XX and 5 XY - from each of the 21 non-SAS 1KGP populations.This was done by iterating through a list of participants sorted by participant ID, and choosing the first 5 XX and 5 XY from each population. As done with the SAS set, these individuals’ reads were then aligned against T2T-CHM13 using bowtie2.4.1, the unaligned and poorly aligned reads were extracted, the reads were assembled using MEGAHIT with default parameters, and filtered for size and contaminants.

### Reference vs reference comparison

Another analysis we performed to highlight the improvements made in T2T-CHM13 over the previous GRCh38 reference is “cross-alignment”. Contigs assembled from unaligned reads against T2T-CHM13 were aligned against those assembled from unaligned reads against GRCh38, and vice versa. We also aligned the unaligned contigs from both references against both references, to demonstrate how T2T-CHM13 is more effective at “rescuing” these contigs. These alignments were done using Minimap2 v2.22 ^45^, and the number of contigs that align or fail to align from and against each reference is noted.

### Categorizing sequence as shared or unique

We evaluated how much of the sequence in the T2T-CHM13 unaligned contigs was unique to the individual it is present in and how much was present in multiple individuals using all-vs-all alignments of the contigs to themselves.

The all-vs-all alignment was performed using Minimap2 v2.22 ^45^ with the strictest “Long assembly to reference mapping” default parameters (asm5), and the resulting output file was used to collapse the shared sequence. To do this, we repeatedly iterate through the output of the all-vs-all alignment and consider contigs with alignment lengths to each other greater than 90% of their own length to be the same sequence. We selected the larger of these sequences to be the representative “super-contig”, and we noted the other sequence as being equivalent to or is derived from this one. We repeated this process until a full iteration through the set occurs without any more sequences being collapsed down, leaving a finalized list of “super-contigs”, a list of which contigs have collapsed into each super-contig, and a list of sequences that have undergone no collapsing and are therefore unique. Combining the list of super-contigs with all non-collapsed sequences gave us our finalized, “non-redundant” set of SAS contigs (**Supplementary Note 8**).

This approach acts as a floor on the amount of shared sequence, as it only merges contigs that are almost entirely identical. However, this approach does not consider contigs that share small subsequences or overlap only slightly at the ends. These situations would be best captured through all-vs-all comparisons and collapsing with no threshold for minimum alignment length. Alternatively, using a similarity metric such as Mash distance ^68^ could quickly estimate the similarity between contigs and decide which ones to merge. Such an approach would account for the case where two contigs have smaller regions of similarity dispersed throughout their length. For Mash distance calculations we used Mash v2.3 ^68^ with default parameters (1000 hashes, sketch and compute overlap in one step) on the Cartesian product of the set of individuals and themselves.

### Assessing non-SAS contribution to SAS contigs

For each of the 210 non-SAS individuals, consisting of 5 XX and 5 XY individuals from the 21 non-SAS 1KGP populations, we aligned their reads using bowtie2 v2.4.1 to the finalized non-redundant set of SAS contigs. We then parsed the alignment output logs to extract their alignment rates, and then further analyzed these counts to generate sub- and super-population summaries.

### Placing assembled sequence

In order to place the contigs against the reference, we followed two approaches: direct alignment of the contigs using Minimap2, or using mate-pair reads that have been mapped to the reference.

For direct alignment, we aligned the contigs against the chosen reference, and reported the locations of all contigs with alignments over >500bp. This alignment and filtering stage was performed using Minimap2 and then a simple post-alignment filtering based on the length of the alignment, which is stored in one column of the output .paf file (**Supplementary Note 3**).

The mate-pair placement method is the approach taken in the construction of the African Pangenome ^21^. During our read extraction step, we explicitly extracted unaligned reads that had mapped mates, and once we knew which contig an unaligned read was assembled into, we could use the mapped mate to anchor the entire contig (**Supplementary Note 3**).

#### Intersections with and analysis of annotated elements

We used the “JHU RefSeqv110 + Liftoff v5.1” annotations generated by the T2T consortium and intersected this with the list of placements generated in the previous step. We computed these intersections using bedtools v2.30.0 ^44^, using the “-wb” parameter to output the details of the intersection annotated element. The bedtools output could then be further analyzed using Python scripts to generate counts for the number of placements per gene, the number of times a gene is intersected with, and other such counts (**Supplementary Note 4**).

We obtained details on the gene types, status, summaries and expressions from their NCBI pages. This can be done using the NCBI E-utilities or simply by scraping the corresponding webpage for each affected gene - we primarily used the latter, due to technical issues with the E-utilities tool. The vast majority of the summary and expression data come from RefSeq and “HPA RNA-seq normal tissues” project ^69^.

#### GWAS intersections with placed contigs

We used the same steps as the annotated region intersections, except this time using the “GWAS v1.0” annotations generated by the T2T consortium. We used “bedtools intersect” with the same configuration to generate the list of all direct overlaps between GWAS elements and placed contigs (**Supplementary Note 4**).

In order to account for placements that are close to but not strictly overlapping the GWAS elements, we implemented a Python script to compare the coordinates of placed contigs against those of known GWAS elements, and compute the distance between all placed contigs in a chromosome and all its GWAS sites. This was done by computing the distance between all GWAS sites and the start and end points of each contig, thus allowing us to both identify all overlaps but also pre-compute all distances between all pairs of contigs and GWAS sites. We could then alter the threshold we want to test for, and count the number of interactions with a distance lower than this threshold.

### Analysis of unplaced sequence

We queried the unplaced sequences against the BLAST “nt” database to ascertain their potential significance, and extracted the top 50 highest scoring hits for each contig. We did this by parsing the BLAST output to extract the lines containing the top matches after each query sequence, before tallying up the number of sequences included in those “top hits”, and outputting these to a table (**Supplementary Note 6**).

### Linkage disequilibrium of shared sequence

From the 13,875 sequences present in multiple individuals, we generated a 640 x 13,875 binary grid representing the presence (1) or absence (0) of each of the shared sequences (or those that have been collapsed down into them) in each of the 640 SAS individuals. Each column in the grid represents the presence or absence of a given contig in each of the 640 individuals in the SAS set, and two contigs that are in LD with each other will have highly similar columns.

We computed a pairwise r^2^ between each of the 13,875 contigs, and then filtered out pairs that had a score higher than our chosen threshold. We then used our placement data to identify pairs of contigs placed within the same chromosome or within a threshold distance, and to identify pairs where one contig is placed and the other is not.

This approach “links” pairs of contigs with high degrees of association; it may be possible to extend this to tie together and place larger groups of contigs or incorporate other data into the association ^33^.

### Pangenome analysis

We tested a range of pangenome aligners, including GraphAligner ^70^, VG ^71^, vg giraffe ^72^, Minigraph ^73^ and Minigraph-Cactus ^74^. We found the vg suites of tools to be easiest to use, but the outputs across the aligners are largely consistent. Regardless of tool, we either forced the output of the aligners to be in the GAM format, or used existing conversion tools to create GAM files. These GAM files can then be converted to the standard SAM/BAM/CRAM format for read extraction (**Supplementary Note 9**). Once in the SAM format we continued to filter and extract reads exactly as we did against the linear references, and repeat the same assembly and filtering steps for these contigs.

Placement was also more complicated due to there being multiple parallel variant sequences, differences in coordinate systems and indexing between aligners (or even different versions of the same aligner), and the increased computational complexity of now performing alignment against a branching sequence graph. We obtained basic coordinates and counts from parsing the various alignment files generated by the different aligners, all of which are run with the finalized list of contigs and the pangenome reference as input and default parameters.

Once placed, there are few resources (GWAS catalogs, annotations, identified significant loci) that have been fully translated into the new coordinates and formats used in these pangenome builds. Therefore, we did not replicate this analysis against the pangenome references.

### Long read validation

We used long read data for 21 SAS individuals from the 1KGP ONT and 20 SAS individuals from Schloissnig et al. in an effort to validate our contigs ^34^. We first construct basic long read assemblies for each of the 21 individuals. This was computed using the Flye assembler ^75^, with default Flye v2.8.1 parameters for “raw-ONT” reads, and with just the long read sets as input (**Supplementary Note 7**).

The previously unaligned short reads were then aligned against these assemblies using bowtie2, and the assembled unaligned read contigs can be aligned using Minimap2 (**Supplementary Note 9**). We used the same tool versions (bowtie2 v2.4.1, and Minimap2 v2.22) and parameters for each of these as the read alignment and contig mapping stages in the analysis of the short read linear reference data.

For short reads, we filtered the resulting SAM alignment file to obtain a list of the reads that now align as well as a list of the reads that remain unaligned. Similarly, the output of the Minimap2 alignment of contigs to this individual-specific long read reference was filtered based on the alignment quality and length.

#### Long read validation of contig placements

We overlapped the list of contigs with high quality alignments in each of the 21 individuals against the list of placed contigs and their intersections with gene and GWAS elements, giving us a list of long read validated intersections. This was done by iterating through the list of placements that intersect an annotated element or GWAS site, and checking if the placed contig had an alignment to the long read assembly.

### RNA-Seq analysis

We used STAR v2.7.10 ^76^ to align the RNA-Seq data obtained from the MAGE dataset ^49^ from each of the 140 individuals against the short read contigs assembled from their unaligned reads against T2T-CHM13, and selected all contigs with large numbers of RNA-Seq alignments (**Supplementary Note 5**).

We tested both single and two-pass alignments, and found little to no changes between these two approaches. Alignment was also tested without splice junctions or annotations, and with a list of known human annotations; once again there was no significant difference. The remaining numerical and memory parameters (including “genomeSAindexNbases” and “limitGenomeGenerateRAM”) were left as default, or amended to STAR’s recommendation (**Supplementary Note 5**).

Once we had identified the most aligned-to contigs, we used the BLAST “nt” database to query for these sequences and report all top hits. As we did for the analysis of unplaced sequence, we parsed the BLAST output for each query sequence and output the top 50 highest scoring hits for each contig (**Supplementary Note 6**).

We also performed an exhaustive all-vs-all alignment of all the RNA-Seq reads across the 140 individuals against all the contigs assembled across the 640 SAS individuals. The parameters for this alignment were slightly adjusted, mainly so that STAR can handle the sharp increase in the number of reads (notably, the “genomeSAindexNbases” and “limitGenomeGenerateRAM” are adjusted as per STAR’s requirements). Similar to the previous analysis, we extracted the most aligned to contigs (>200 RNA-Seq alignments per contig), and ran them through the same BLAST query process. The top 50 BLAST hits (if there are that many) for each contig were then extracted.

### Biobank analysis

We used the UK Biobank Phenotype Manifest from the Pan-UK Biobank effort ^53^ to obtain our chosen set of 21 quantitative biomarker traits. Using the provided data for each trait, we were able to extract a list of all chosen loci in the genome and their potential association with each of the traits. The manifest represents this association as the negative log of the p-value, and this p-value is computed for both the entire UK Biobank set and for each of the individual ancestry groups within the Biobank. For our analysis, we used the p-values across the “CSA” ancestry group and the entire UK Biobank set. The Genes and Health dataset ^52^ is formatted in a similar way, and contains the associations between all chosen loci in the genome and 42 quantitative biomarker traits.

For each trait, we then “swept” across all potential loci, and identified all loci that have significance above our chosen p-value threshold. For all significant loci within 50 kbp of each other we selected the highest scoring loci (i.e. the one with the most significant p-value) as the representative “peak” for the entire group. We computed the pairwise distance between all selected peaks for a given trait and all placed contigs across the SAS set, and identified all peak/placed contig pairs that are within 1 Mbp of each other. This process is repeated for all 21 biomarker traits in the UK Biobank, and for the 42 quantitative biomarker traits in the Genes and Health dataset.

### Comparison to existing variant discovery tools

We compared the assembled contigs from 21 SAS 1KGP individuals with LR data to the variant calls made in those individuals using Manta ^46^ and Lumpy ^47^. Both tools were run with default parameters using the output of the alignment stage (**Supplementary Note 10**). Manta uses the .cram file from the alignment stage directly, while there is an initial preprocessing step to extract the split and discordant reads for Lumpy.

Both tools output a vcf file that detailed the structural variants they find, which we parsed to determine the location, type (“INS” for insertions, “BND” for breakends of large insertions in Lumpy) and size of the SVs. We then compared this against the list of contigs assembled from these individuals, the contig placements in this set, and which of the contigs were validated with long reads.

For PopIns2 ^48^, we selected 10 samples from the 1KGP SAS set, and followed the steps outlined in the PopIns2 Github repository using default parameters, starting from the alignments of these individuals’ reads against T2T-CHM13 (**Supplementary Note 10**). We then parsed the resulting assembly and vcf files to compare against the assembled and placed sequence from our approach.

### Data and Code Availability

Our analysis was performed on both our local computing cluster at JHU and on the NHGRI AnVIL cloud platform, with the initial experiments performed on the former and the full set being analyzed on the latter. Experiments on AnVIL were run with a number of Docker environments and using WDL workflows.

The analyses were computed using a range of established bioinformatics tools and custom scripts, and plotting was performed using existing Python libraries, such as sklearn ^77^ or matplotlib ^78^. Each tool used has been mentioned in the relevant methods section, and all relevant scripts and documentation can be found at github.com/arun96/SouthAsianGenomeDiversity. The AnVIL workspace used for this analysis can be found at https://anvil.terra.bio/#workspaces/anvil-dash-research/south-asian-genome. The data generated by this work can be found on Zenodo, at doi.org/10.5281/zenodo.17419004; this includes the final collapsed contigs across the SAS set, details about the placed locations of contigs, the Repeatmasker outputs, and several other pieces of information.

## Supporting information

Supplementary Results, Methods, Figures and Notes

## Declarations

### Author contributions

A.D. was the primary author of this work, designed and ran the experiments, wrote the manuscript and designed the figures, all under the supervision of M.C.S. M.C.S, R.C.M. and A.B. advised throughout the project, and helped with writing and editing the manuscript and figures. All authors read and approved the final manuscript.

## Acknowledgements

We would like to thank Dr. Dylan Taylor for providing us with early access to the RNA-Seq data, and to the members of the 1KGP ONT consortium for providing early access to the long read data used for validation. This work was supported by NIH awards U24HG010263, R03CA272952, U01HG013744, and R35GM133747.

