## Supplementary Results, Methods, Figures and Notes for "Assembling unmapped reads reveals hidden variation in South Asian genomes"

### Supplementary Materials

#### Supplementary Results

##### Alignment and assembly of poorly aligned reads

In addition to extracting unaligned reads against the references, we also investigated the looser threshold of poorly aligned reads, defined as reads that align with quality score <20, which results in about 20-30x as many extracted reads as the unaligned metric. As with the unaligned reads, there was a decrease in the number of poorly aligned reads when moving to T2T-CHM13; however the extent of this decline varies greatly across samples, ranging from 11% to 42%.

Assembling the poorly aligned reads against T2T-CHM13 resulted in 50 Mb of sequence assembled per individual, filtered down to 20 Mb in non-contaminant >1 Kb contigs. Assembling the poorly aligned reads against GRCh38 resulted in >60 Mb of sequence in assembled, filtered contigs, approximately 3x what was assembled against T2T-CHM13 (**Supplementary Figure 16**). Against Minigraph and Minigraph-Cactus, we observe decreases in the amount of sequence assembled from these poorly aligned reads, down to 15-18 Mb per individual in non-contaminant >1 Kb contigs.

##### Comparison to non-SAS placements

We repeated the placement and intersection steps with unaligned read contigs from the 210 non-SAS individuals from the 21 non-SAS populations in 1KGP. Across this set there were 4,105 intersections with 88 unique non-LOC genes. The distribution of intersections across the genome was similar to what we see with the SAS set, and most of the intersected genes were intersected in only a few individuals (**Supplementary Figure 26**).

Of the 88 intersected non-LOC genes, 70 were also intersected in the SAS set's placements. There were 66 genes that only have intersections in the SAS set, and 18 that were only found in non-SAS individuals (**Supplementary Figure 27**). The majority of genes intersected in both sets were intersected in 10 or fewer individuals, with 19 intersected in >100 and 7 intersected in >500.

55 of the genes present only in the SAS set were protein coding, and all 55 were intersected in <2% of the set. They included a number of genes associated with ocular disorders and conditions (CPAMD8, KIF4, MYO7A, PPEF1, WDR17), and genes associated with forms of cancer (ITGA9, KIF4, PTPRO, SPECC1). 15 of the genes present only in the non-SAS set were protein coding, and all but two were intersected in <3 individuals. The two most prevalent genes are IYD and PTPRN2, which are intersected in >25 individuals, and mutations in which are associated with congenital hypothyroidism and insulin secretion respectively.

When grouped by the 1KGP super-population groups, the SAS populations have a similar number of average gene intersections per individual as the AMR and EUS groups, and slighter lower number than the AFR groups, and slightly higher counts than the EAS groups (**Figure 3d**, **Supplementary Figure 28**).

##### GWAS intersections with the non-SAS set

We repeated this GWAS catalog analysis with the placed contigs from the non-SAS set. At a threshold of 100 bases, there were 2,240 potential interactions across 26 unique GWAS sites, rising to 5,190 across 90 unique sites and 19,268 across 551 unique sites at thresholds of 5 Kb and 10 Kb respectively. As the non-SAS set is roughly a third of the size of the SAS set, the total number of potential interactions was what we expect. However, the number of unique GWAS sites that are within the threshold distances of a contig did not decrease proportionally in the non-SAS set.

##### Identifying patterns within the shared contig set

We investigated if the sharing of certain sequences could allow us to identify patterns within the SAS set. This was done by recording the presence/absence of each of the ~14K shared sequences in each of the 640 individuals, and performing PCA on the resulting matrix.

We found the 1KGP SAS individuals split into two groups (split approximately 5:3), with the 39 SGDP individuals clustering separately. This ratio loosely resembled what we would expect from the Hardy–Weinberg principle, with $p = 0.75$ and $q = 0.25$, suggesting that these clusterings may be due to the presence or absence of a particular allele in each of the sub-groups. These clusters became more clear when we limited the shared sequences to only those present in >5 individuals, and further to include sequences present in both the 1KGP and SGDP sets. With these additional requirements, the SGDP individuals still clustered separately, while the 1KGP individuals split into two main clusters (**Supplementary Figure 29, Supplementary Figure 30**).

### Supplementary Methods

#### Alignment and assembly of poorly aligned reads

We tested a number of definitions for what defines a poorly aligned reads, including a range of quality score thresholds and the edit distance of a read alignment. We settled on selection based on a quality value below q20, as this was the best compromise between capturing potentially interesting reads without broadening the set too much.

#### PCA clustering based on presence/absence of shared sequence

We performed PCA on the scaled 640 x 13,875 binary grid representing the presence (1) or absence (0) of each of the 13,875 shared sequences (or those that have been collapsed down into them) in each of the 640 SAS individuals. As we added further constraints, we selected the relevant columns that correspond to those sequences. For example, when narrowing down to only the 660 contigs that are shared in >5 individuals, placed again T2T-CHM13, and present in individuals in both the SGDP and 1KGP, we narrowed the grid down to 640 x 660, and performed PCA on the resulting grid.

We then plotted each of the first 10 PCs against each other. To investigate the loadings of each contig, we multiplied each component by the square root of its corresponding eigenvalue. We then plotted the loading for each contig, sorted across the genome. All PCA analysis and plotting was performed in Python using sklearn and matplotlib.

### Supplementary Figures

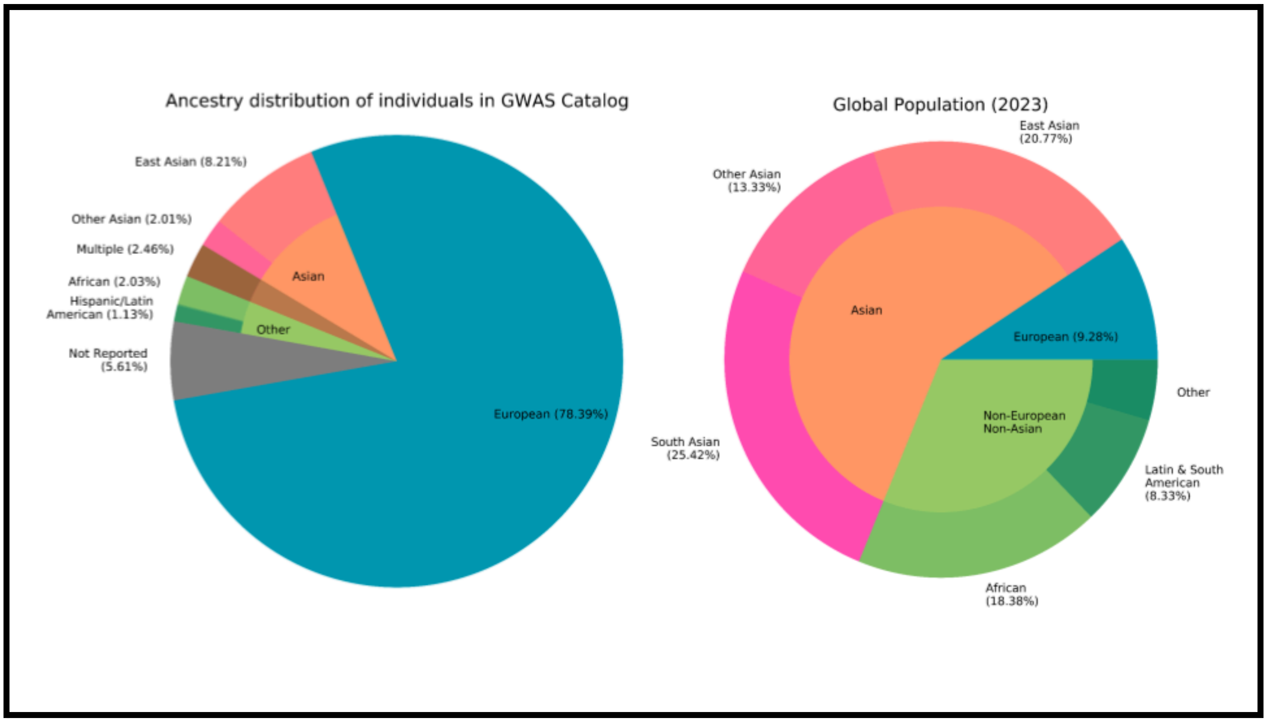


**Supplementary Figure 1: Discrepancies in representation in genomics.** The disproportionate presence European populations have in genomics, with South Asians and Africans having the least representation; GWAS individual ancestry data is from Sirugo et. al. (1) using the GWAS catalog from EBI through January 2019, the population data is from UNFPA (25) as of 2023.

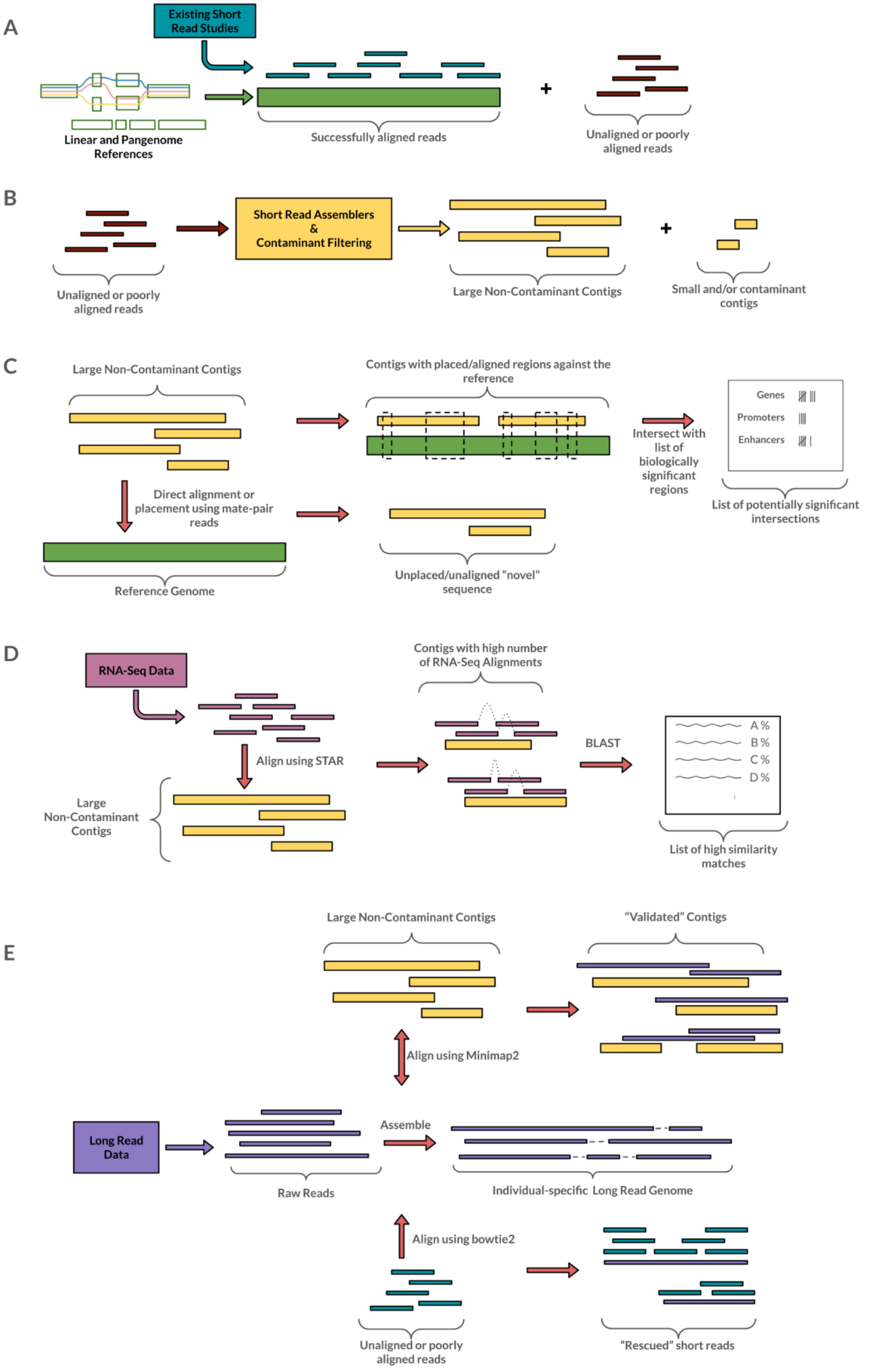
A
Existing Short Read Studies
Linear and Pangenome References
Successfully aligned reads
+
Unaligned or poorly aligned reads
B
Short Read Assemblers & Contaminant Filtering
Unaligned or poorly aligned reads
Large Non-Contaminant Contigs
Small and/or contaminant contigs
C
Large Non-Contaminant Contigs
Contigs with placed/aligned regions against the reference
Intersect with list of biologically significant regions
Genes
Promoters
Enhancers
List of potentially significant intersections
Direct alignment or placement using mate-pair reads
Reference Genome
Unplaced/unaligned "novel" sequence
D
RNA-Seq Data
Align using STAR
Large Non-Contaminant Contigs
Contigs with high number of RNA-Seq Alignments
BLAST
A %
B %
C %
D %
List of high similarity matches
E
Large Non-Contaminant Contigs
"Validated" Contigs
Align using Minimap2
Long Read Data
Raw Reads
Assemble
Individual-specific Long Read Genome
Align using bowtie2
Unaligned or poorly aligned reads
"Rescued" short reads

**Supplementary Figure 2: Pipeline Overview.** Figures **(a)** through **(e)** outline the key steps of our method: **(a)** We align existing short read data against a range of chosen reference genomes, and extract unaligned or poorly aligned reads. **(b)** These extracted reads are assembled into larger contigs, with smaller or contaminant contigs discarded during the filtering process. **(c)** These contigs are then compared against the reference again, with some being placed and their placements being investigated, and others being identified as novel or highly variant sequences. **(d)** We can evaluate the functional impact of these contigs using auxiliary RNA-Seq data, and then use BLAST to search for contigs with a large number of RNA-Seq matches. **(e)** We use long read data from a subset of these individuals, and simple long read genome assemblies of each individual's reads, to perform validation of the assembled unaligned read contigs and to rescue previously unaligned short reads.

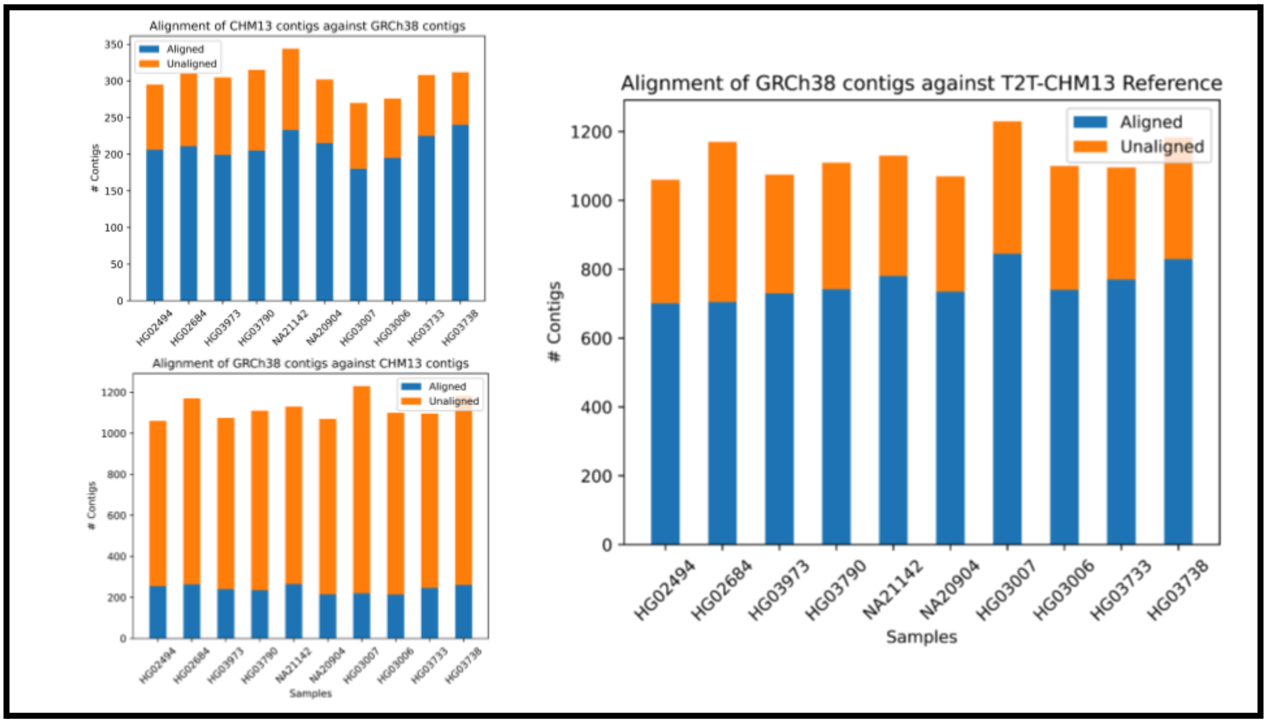


**Supplementary Figure 3: T2T-CHM13 vs GRCh38 cross-alignment.** Cross-alignment results for the T2T-CHM13-relative unaligned contigs and the GRCh38-relative unaligned contigs. Majority of T2T-CHM13 contigs align well to the GRCh38 contigs (**top left**), but the majority of the GRCh38 contigs do not align well back (**bottom left**). This is to be expected, as T2T-CHM13 resolves a number of issues and gaps present in GRCh38, which is where a number of the GRCh38 contigs are assembled from. However, the majority of GRCh38 contigs have a good alignment to the T2T-CHM13 reference itself (**right**), further confirming that these were artifacts or unresolved sequences from GRCh38 that have been now resolved in T2T-CHM13.

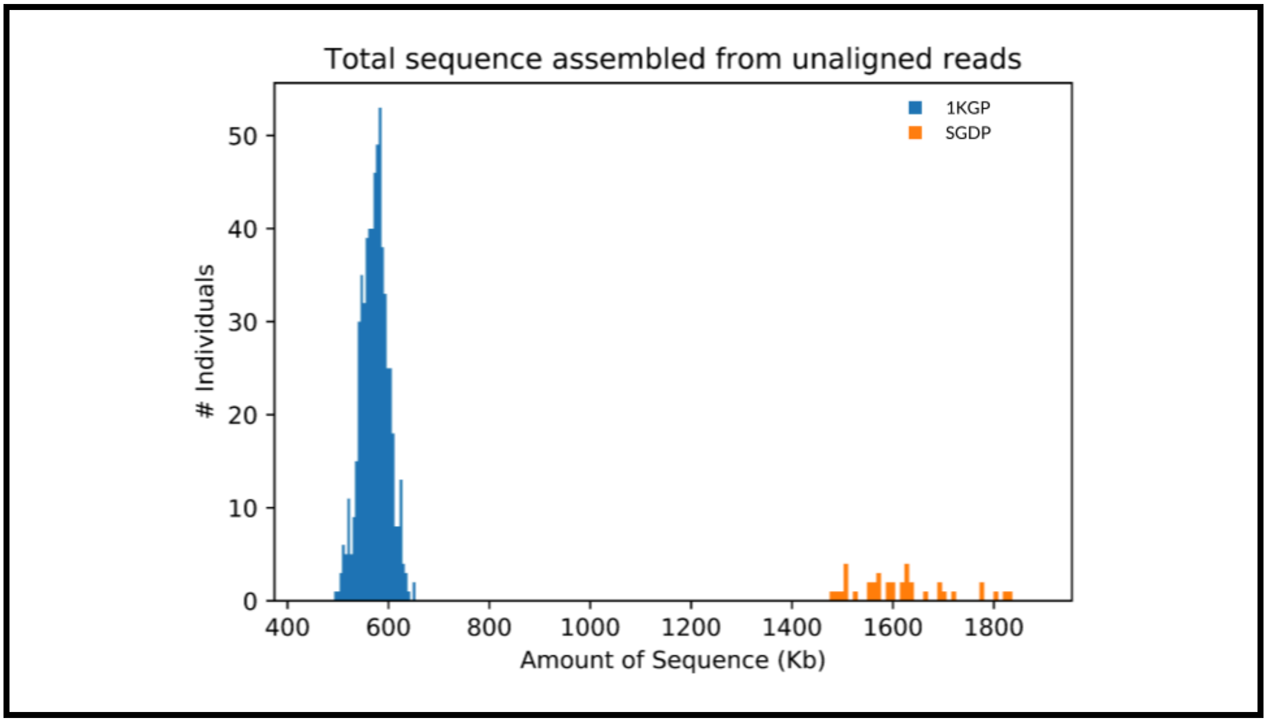


**Supplementary Figure 4: 1KGP vs SGDP assembled sequence distribution.** Comparison of the amount of sequence assembled from unaligned reads into >1 Kb non-contaminant contigs in the 1KGP set (blue) and the SGDP set (orange). Unaligned reads from the SGDP set assemble into, on average, 3-4x as much sequence per individual.

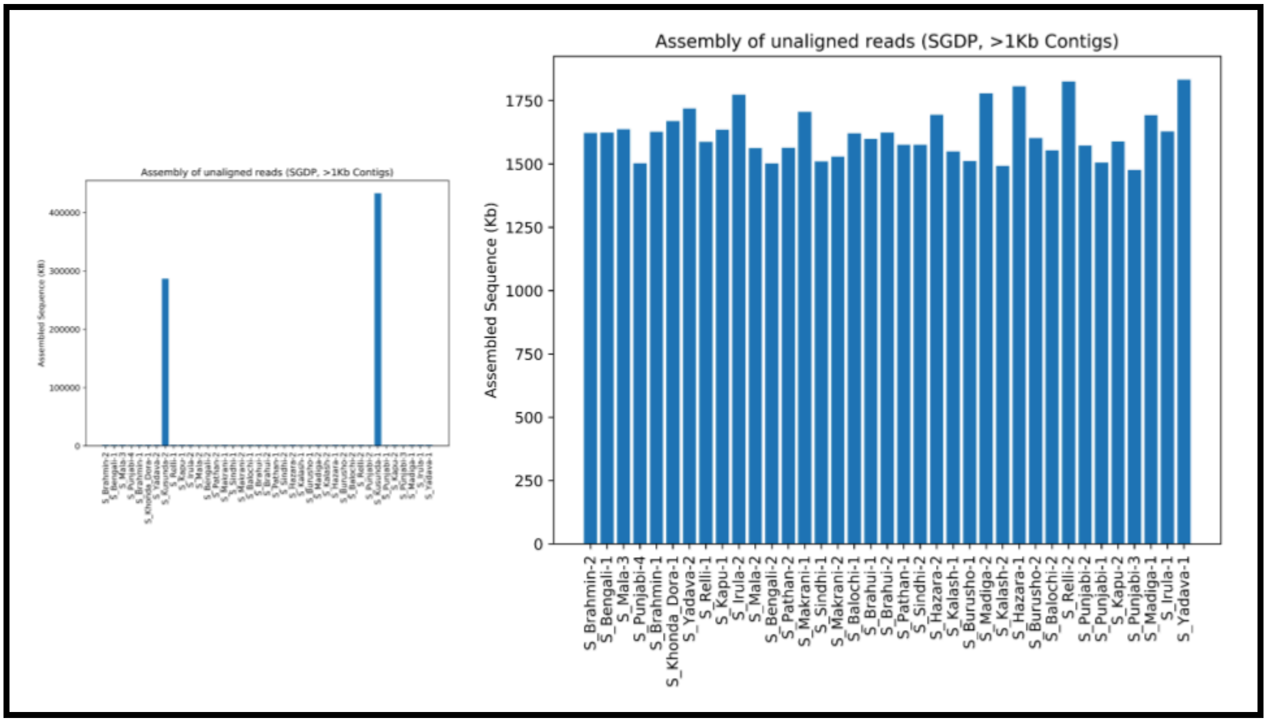


**Supplementary Figure 5: SGDP total assembled sequence:** Counts of total assembled sequence from unaligned reads against T2T-CHM13 across the 39 SGDP SAS individuals. **(a)** Distribution with the two outlier Kusunda individuals included, and **(b)** with these outliers removed.

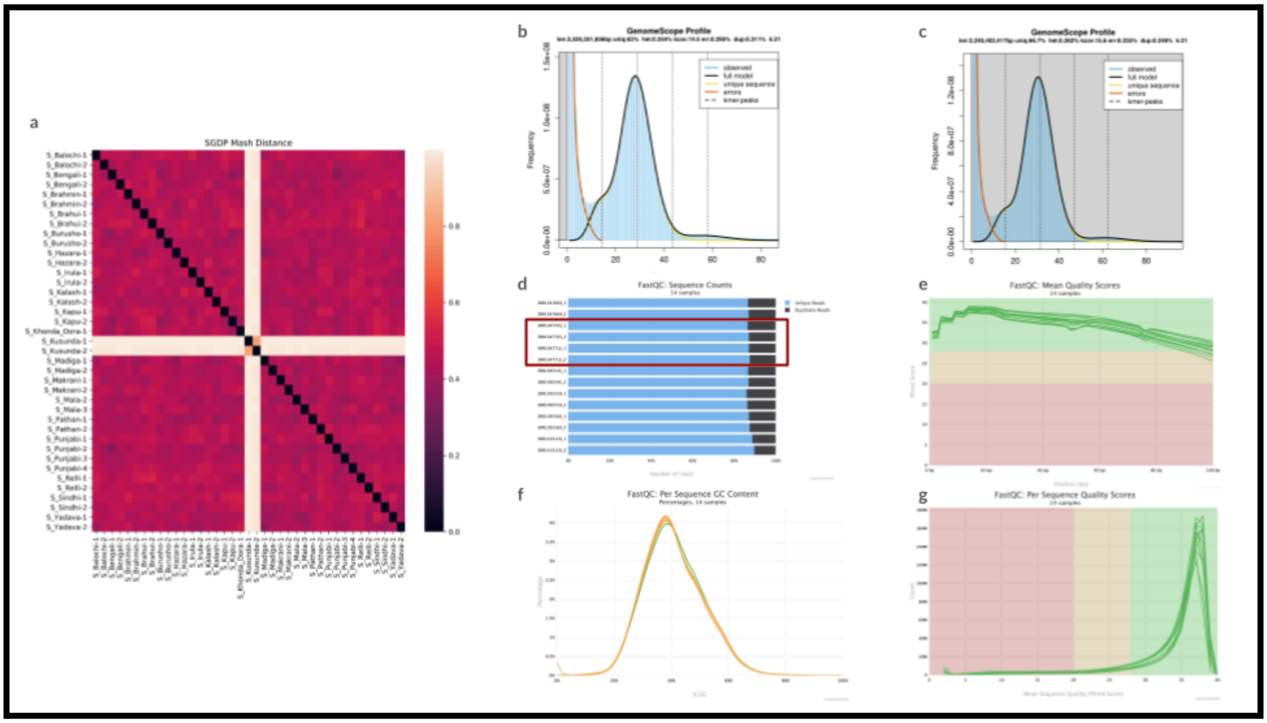


**Supplementary Figure 6**: **Analysis of Kusunda Outliers**. The two Kusunda individuals from the SGDP set assemble 100x as much sequence from their unaligned reads than any other individual in the SGDP set, in contigs as large as 700 Kb. **(a)** The assembled sequence in these two individuals is highly similar, and dissimilar to the sequence assembled from unaligned reads in the other 37 individuals, as shown in this Mash similarity visualization (a clustering based on this distance can be found in **Supplementary Figure 25**). **(b)** & **(c)** Genomescope plots generated from the input reads of the two Kusunda individuals, which show the estimated genome size to be >3.5Gb, compared to the 2.9Gb estimates for the rest of the SAS set. **(d)** - **(g)** Key metrics from FASTQC of the four read sets of the two Kusunda individuals and ten read sets of five more SGDP individuals. The number of unique reads **(d)**, mean **(e)** and per sequence **(g)** quality scores and GC content **(f)** are consistent across the seven individuals. BLAST analysis also does not obviously flag any of this sequence as from a potential contaminant.

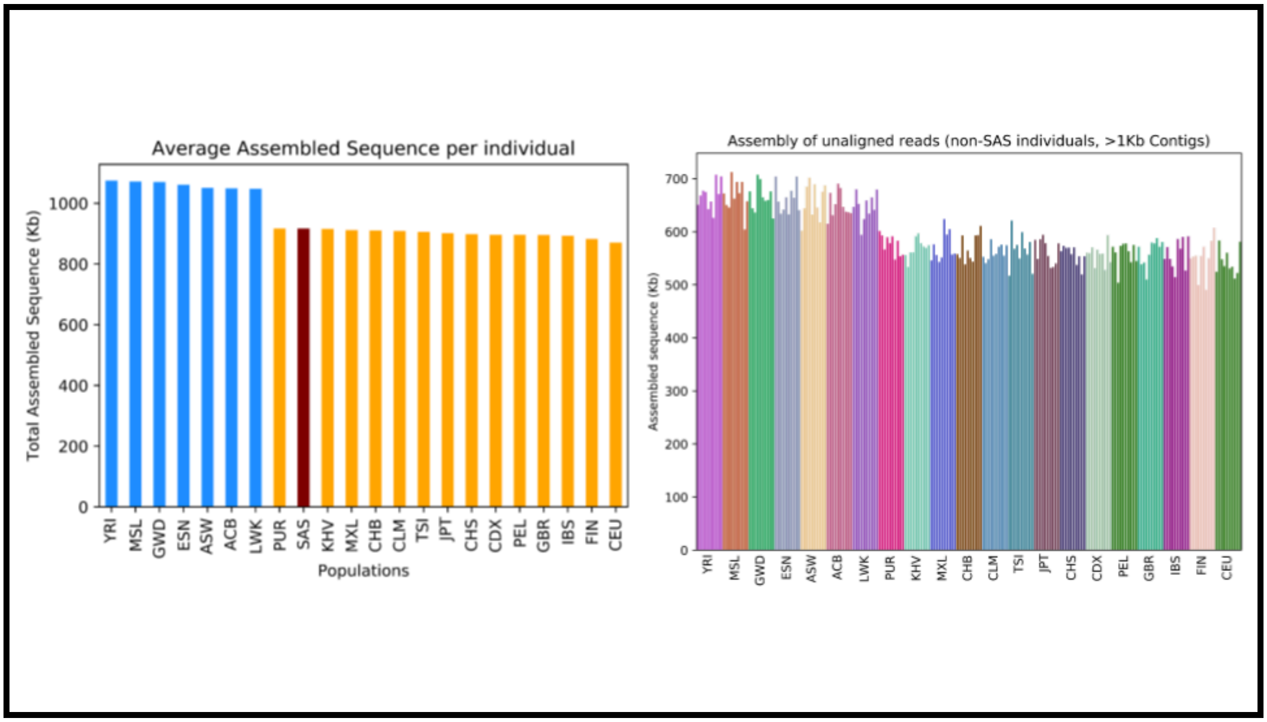


**Supplementary Figure 7: Population-level sequence comparison. (Left)** Comparison of the amount of sequence assembled from unaligned reads across 26 1KGP populations, sorted by the average amount of assembled sequence per individual in the population. The 21 non-SAS populations consist of 5 XX and 5 XY individuals each, with the SAS superpopulation (red) containing all 600 SAS 1KGP individuals across the 5 sub-populations. The 7 AFR populations are in blue, and the 14 non-SAS, non-AFR populations are in orange. **(Right)** The amount of sequence present in each of the 10 individuals in each of the 21 non-SAS populations, sorted by the average amount of assembled sequence in each population and colored based on which of the 21 non-SAS populations they occur in. The first 7 populations (YRI-LWK) are from the AFR superpopulation, while the PUR subpopulation is the only non-AFR group with a higher amount of assembled sequence on average than the average across the 1KGP SAS set.

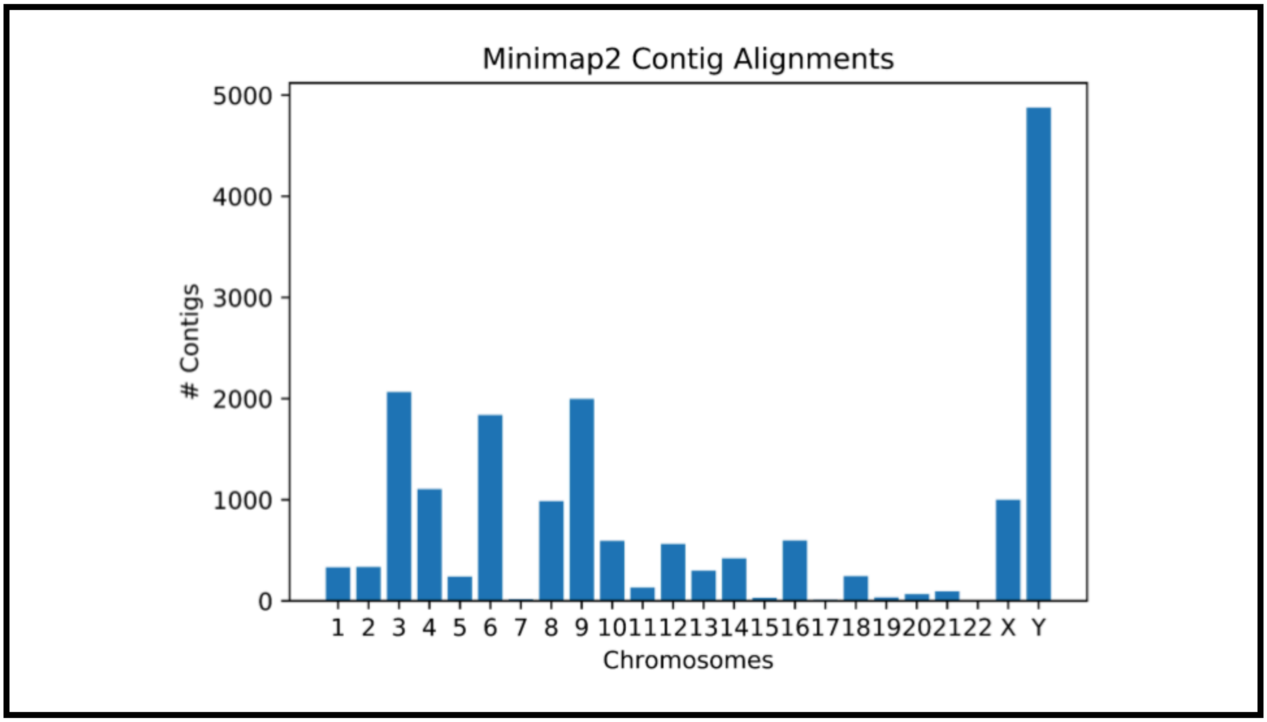


**Supplementary Figure 8: Minimap2 contig alignments.** Number of contigs placed by direct alignment in each of the 24 chromosomes. For these placements, we require a > 500 bp stretch of the contig to be aligned well to the reference genome.

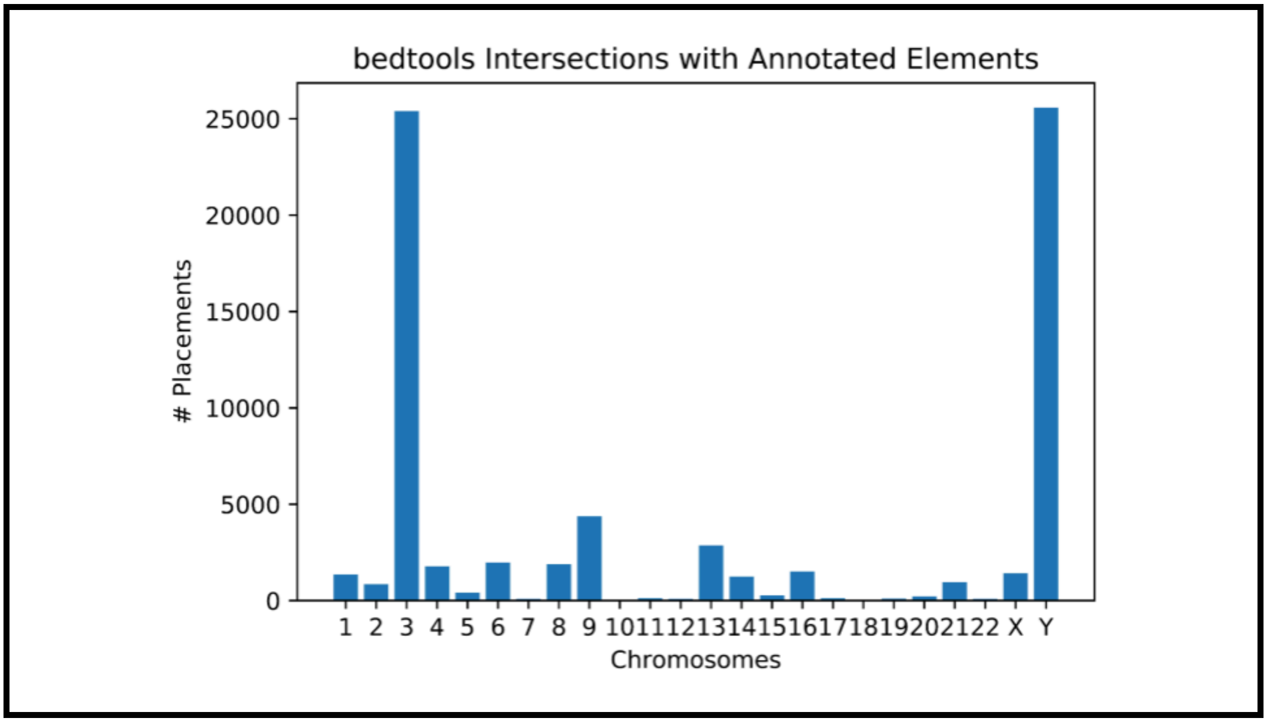


**Supplementary Figure 9: Bedtools intersections with annotated elements.** Number of intersections between the placed contigs and annotated elements in each of the 24 chromosomes.

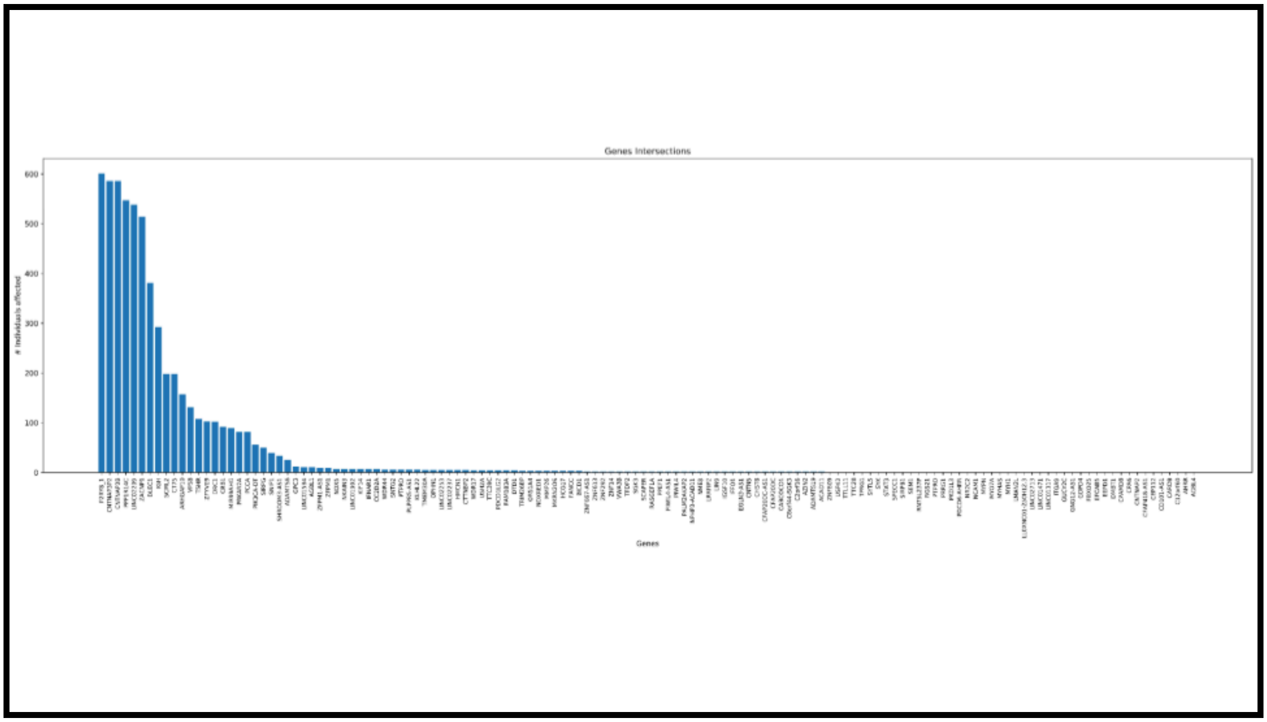


**Supplementary Figure 10: SAS gene intersection distribution.** Number of individuals each of the intersected non-LOC genes are intersected in. Full details and counts are in **Supplementary Table 1**.

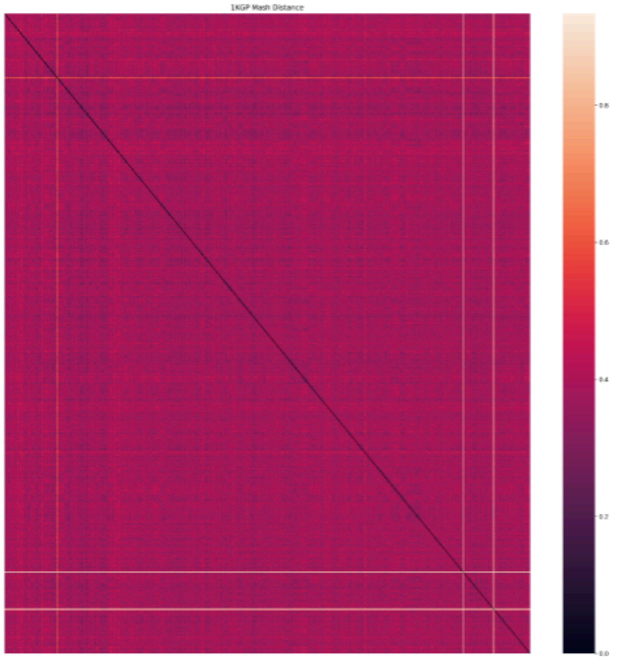


**Supplementary Figure 11: 1KGP Mash Distance.** Mash Distance between the assembled unaligned read contigs from the 601 1KGP SAS individuals. The distances range between 40-50% for the whole set, with a handful of outliers.

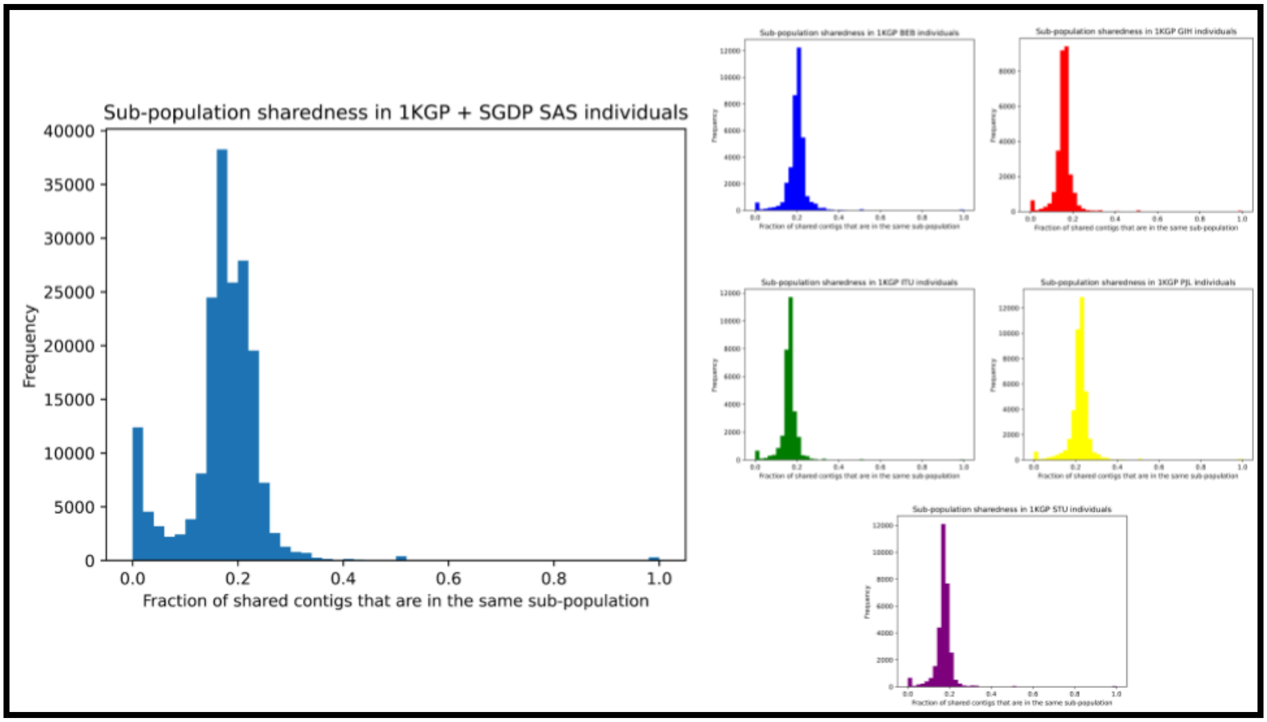


**Supplementary Figure 12: Sub-population contig sharedness.** Fraction of the shared contigs that are shared between individuals in the same sub-population, both across the whole set (**left**), and in each of the five SAS 1KGP populations (**right**). These plots are generated by iterating through every assembled contig, and keeping track of the number of contigs it was similar to during the all-vs-all alignment phase. For each contig, we then track the number of "similar contigs" that were also in the same population as the individual it was assembled from. As each of the 5 1KGP populations make up ~20% of the set each, if variation was spread evenly across the entire SAS cohort, we expect ~20% of the similar contigs to be from the same group. If significant amounts of variation were present only in a single population, the fraction would be higher for contigs of that group, and the peak would shift to the right for that plot. In the overall plot, the fact that the SGDP individuals only have 1-3 members of their community in the set means they all fall into the left-most bin, creating a small second peak absent when looking at each 1KGP population.

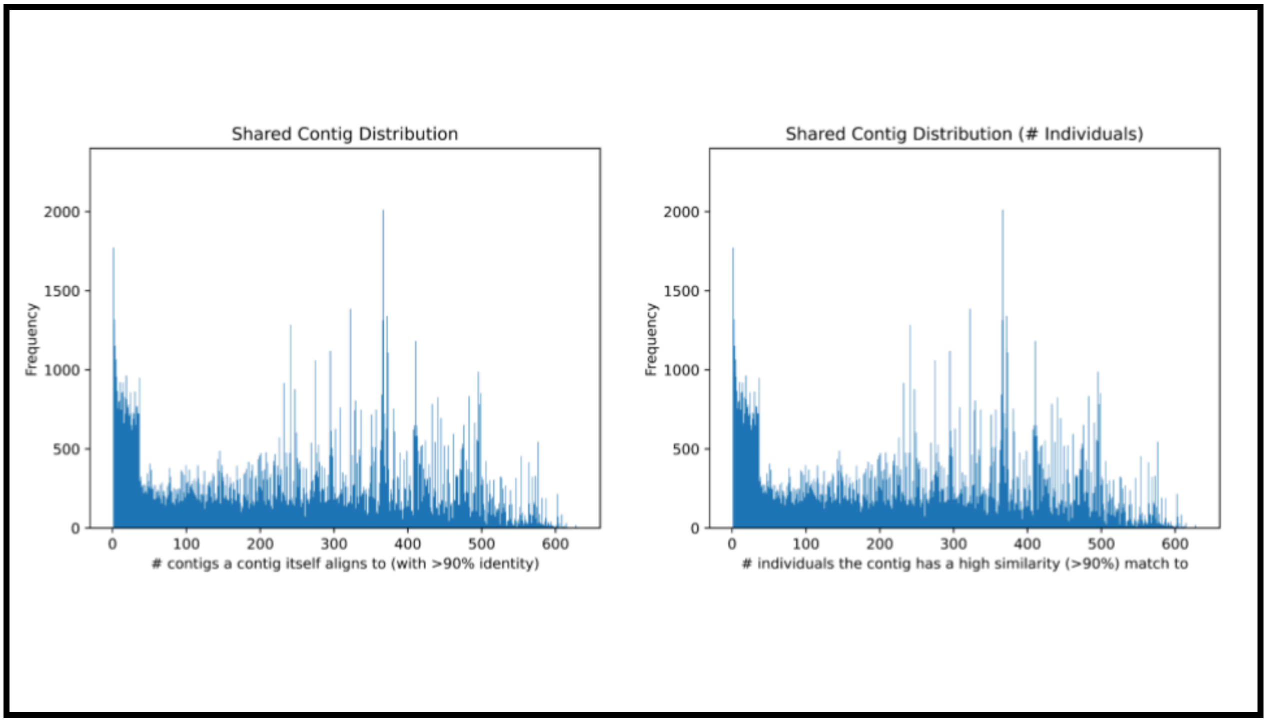


**Supplementary Figure 13: Shared contig distribution.** Distribution of the number of contigs (**left**) and individuals (**right**) each of the shared contigs have similarity to.

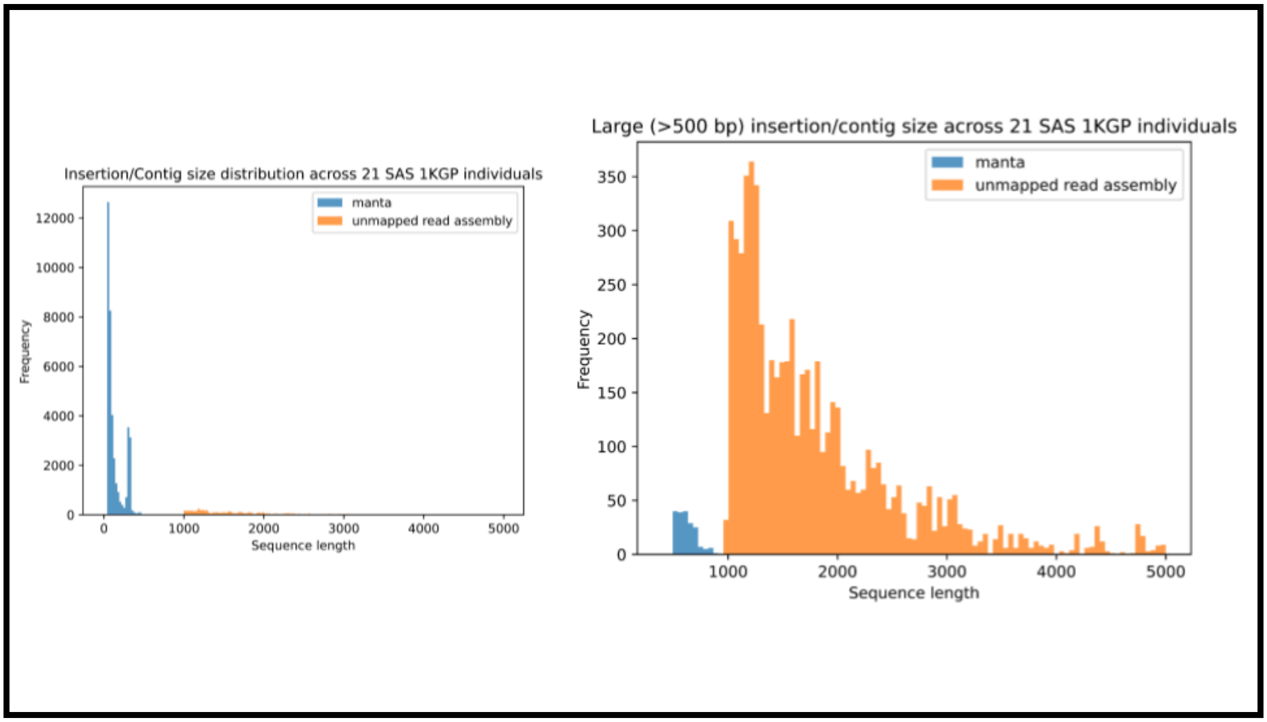


**Supplementary Figure 14**: **Comparison of assembled sequence to Manta insertions**. Manta largely detects small insertions (**left**), with significantly fewer insertions larger than 500 bp being called, and none above 1 Kb (**right**). In comparison, our approach focuses only on > 1 Kb contigs assembled from unaligned reads, resulting in many times as much sequence being discovered.

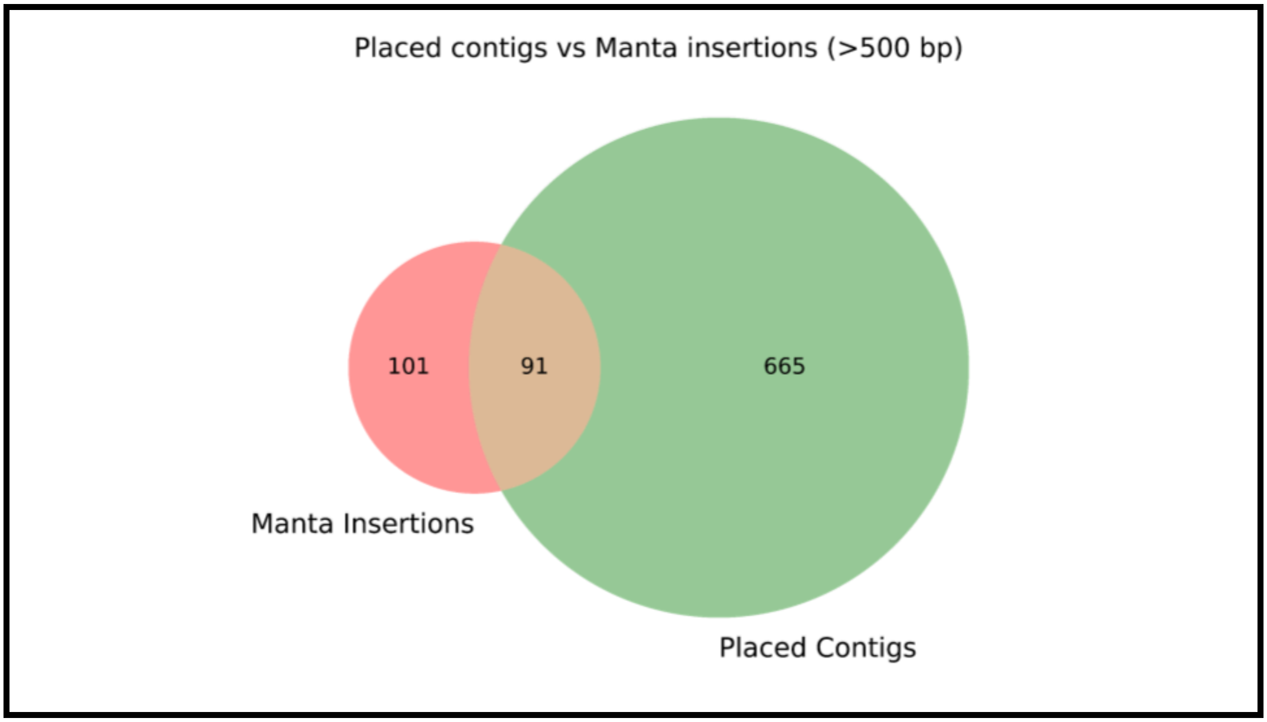


**Supplementary Figure 15**: **Comparison of placed contigs to Manta insertions**. Comparison of the number of large (>500 bp) insertions called by Manta against the number of placed, >1 Kbp contigs we find through the assembly of unmapped reads.

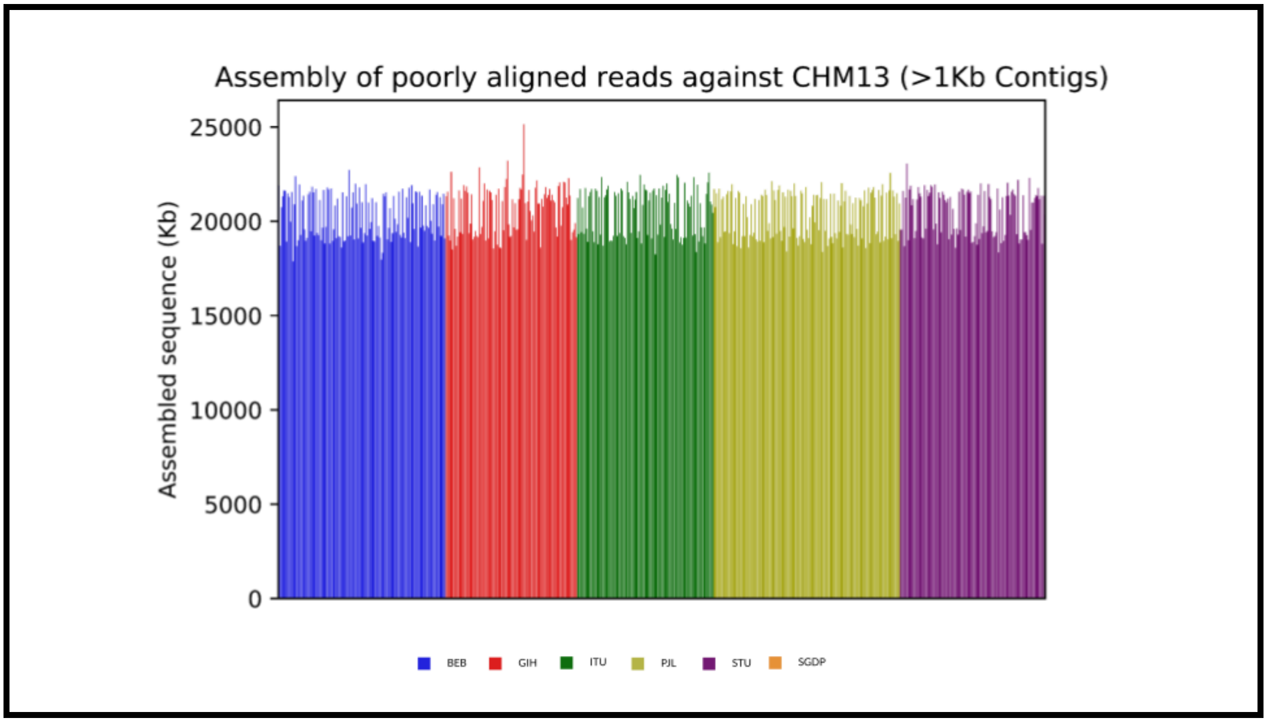


**Supplementary Figure 16: Poorly aligned read sequence assembly against T2T-CHM13.** Amount of assembled sequence from poorly aligned (<q20) reads in >1 Kb, non-contaminant contigs per individual across the 1KGP set.

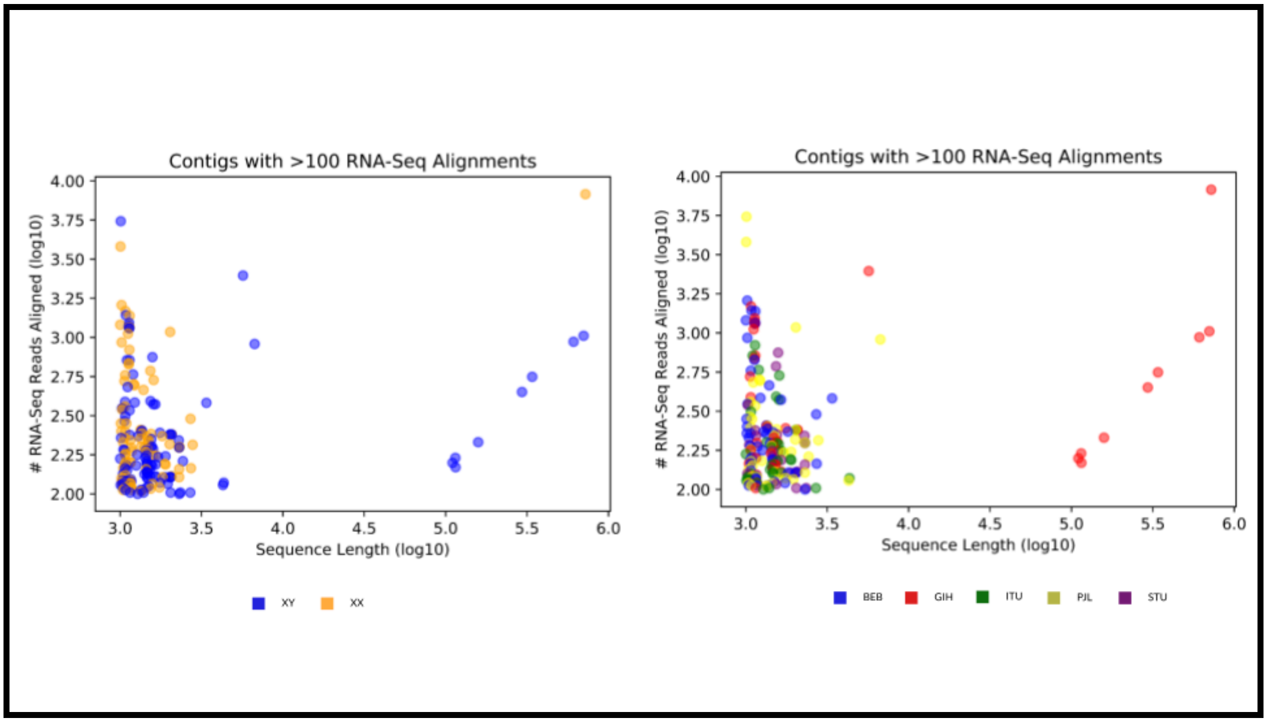


**Supplementary Figure 17: Lengths of 1KGP RNA-Seq aligned contigs.** All contigs from the 1KGP set that have >100 RNA-Seq alignments against the 140 individual RNA-Seq dataset, with the number of RNA-seq alignments plotted against the contig length, stratified based on karyotype **(left)** and sub-population **(right)**.

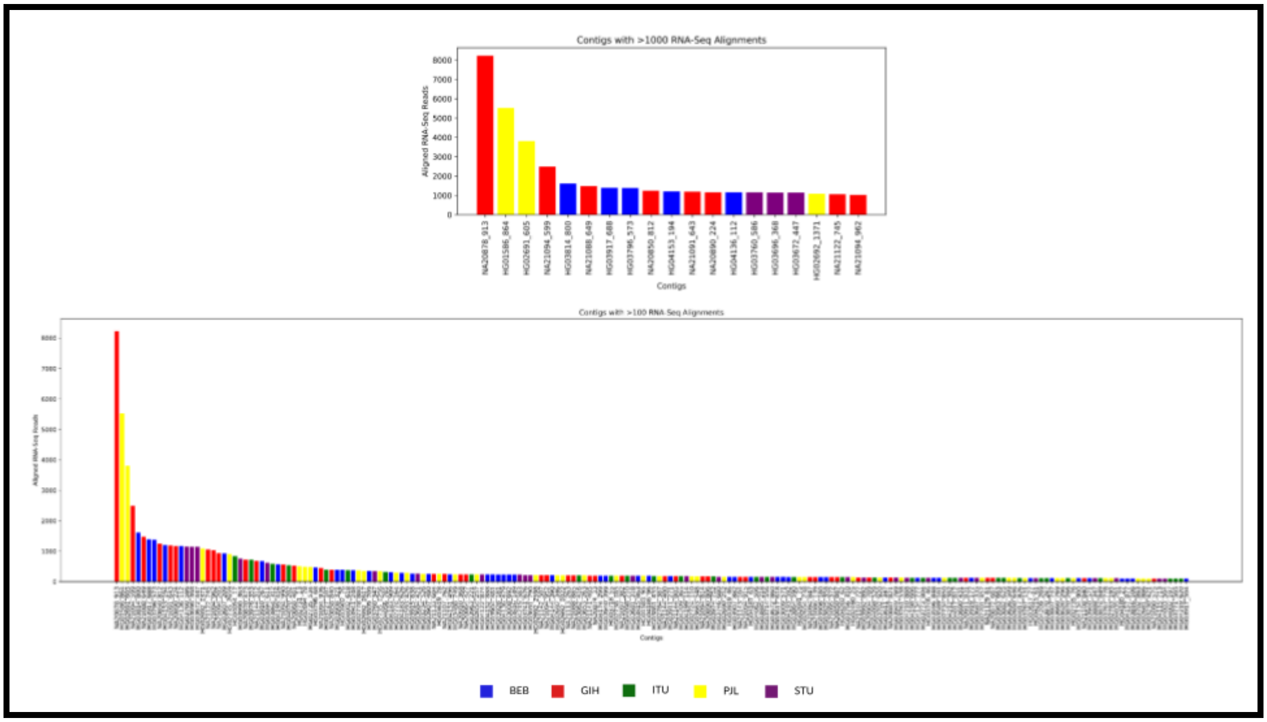


**Supplementary Figure 18: High-scoring RNA-seq aligned contigs.** All contigs from the 1KGP set that have >1000 **(top)** or >100 **(bottom)** RNA-Seq alignments against the 140 individual RNA-Seq dataset, stratified based on sub-population. The BLAST results of these contigs can be found in **Supplementary Tables 3 and 4**.

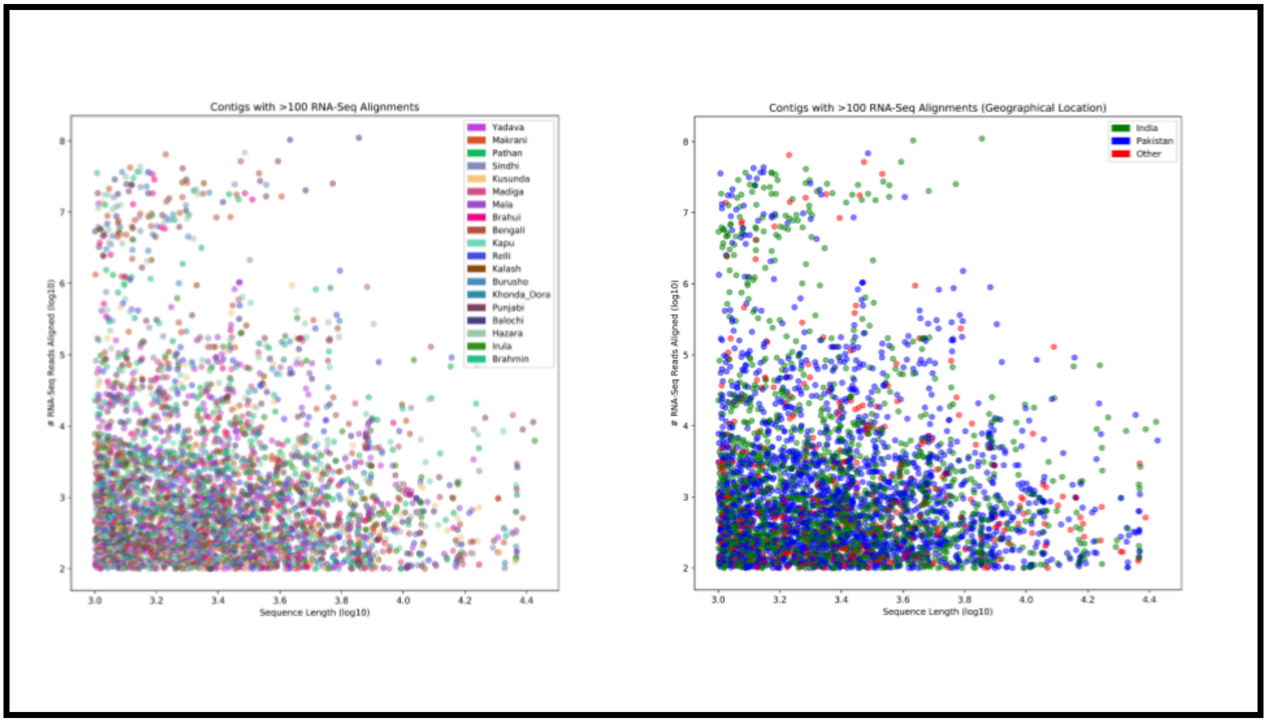


**Supplementary Figure 19: Lengths of SGDP RNA-Seq aligned contigs.** All contigs from the SGDP set that have >100 RNA-Seq alignments against the 140 individual RNA-Seq dataset, stratified based on sub-population **(left)** and geographical location **(right)**. The BLAST results of these contigs can be found in **Supplementary Table 3**.

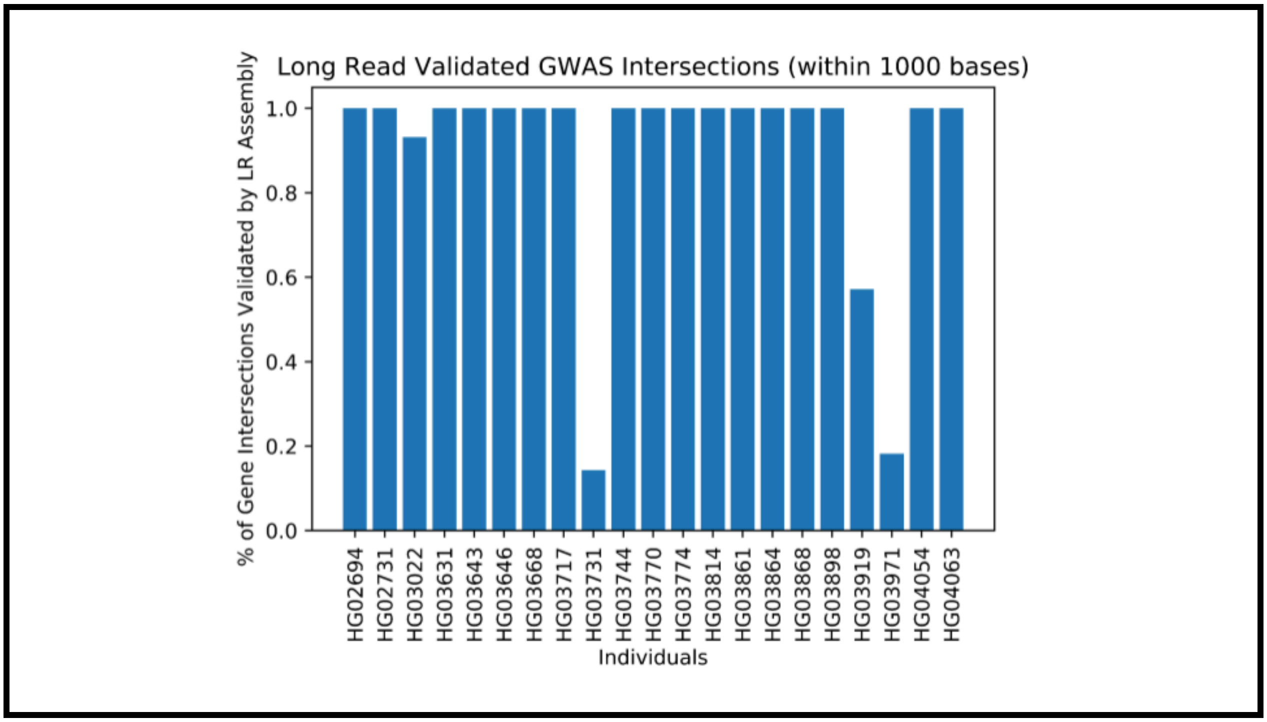


**Supplementary Figure 20: Long read validation of GWAS interactions.** Fraction of GWAS interactions (placements within 1 Kb of a GWAS site) in 21 individuals with long read data that are in “long read validated” contigs.

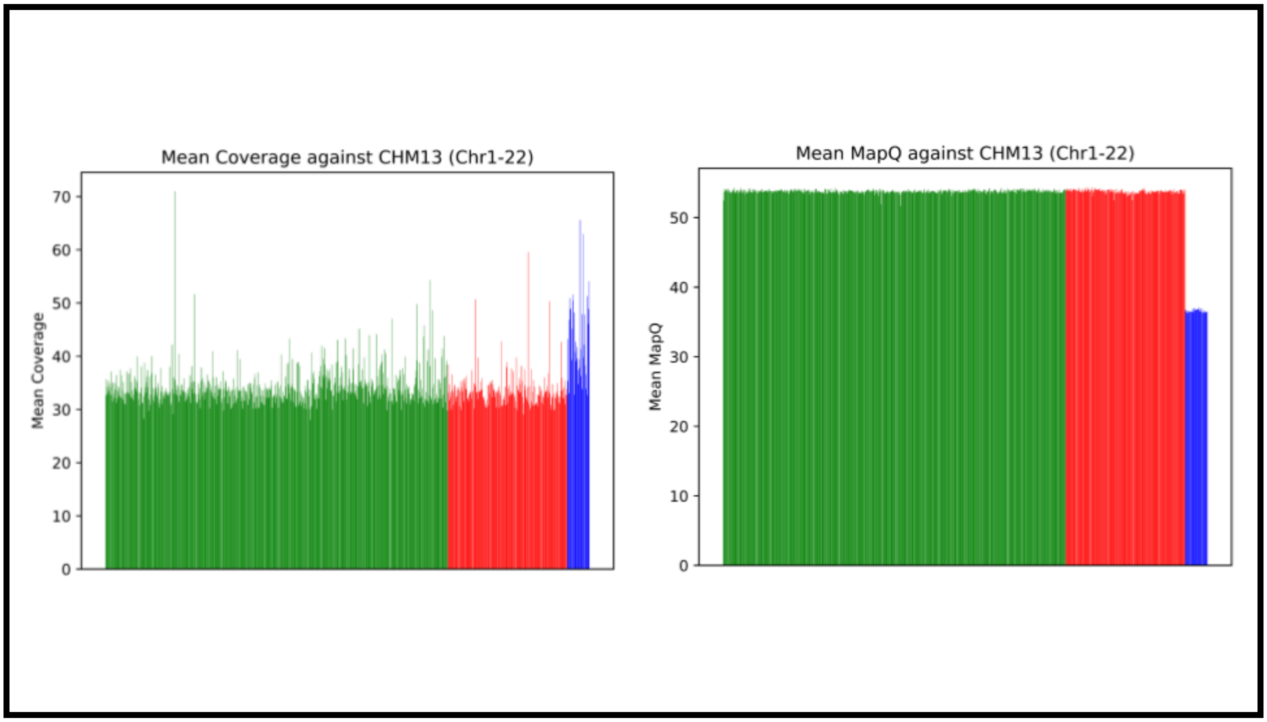


**Supplementary Figure 21: Average coverage and mapping quality against T2T-CHM13.** Mean coverage (**left**) and mapping quality (**right**) of the 1KGP SAS (green), 1KGP non-SAS (red) and SGDP (blue) reads against the T2T-CHM13 autosomal chromosomes.

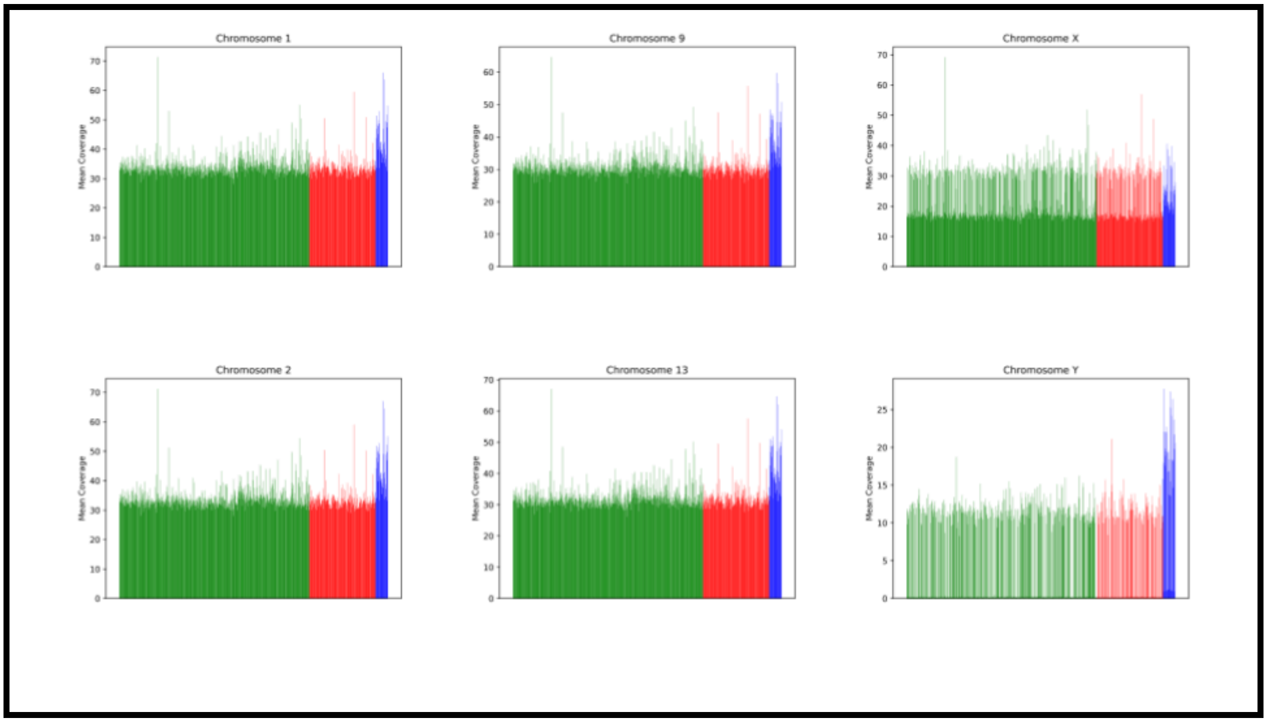


**Supplementary Figure 22: Read coverage depth against T2T-CHM13.** Mean read coverage depth for alignments against T2T-CHM13 across the 1KGP SAS (green), non-SAS (red) and SGDP SAS (blue sets) for chromosomes 1, 2, 9, 13, X and Y. Across all 24 autosomal and sex chromosomes, the SGDP individuals have higher average coverage.

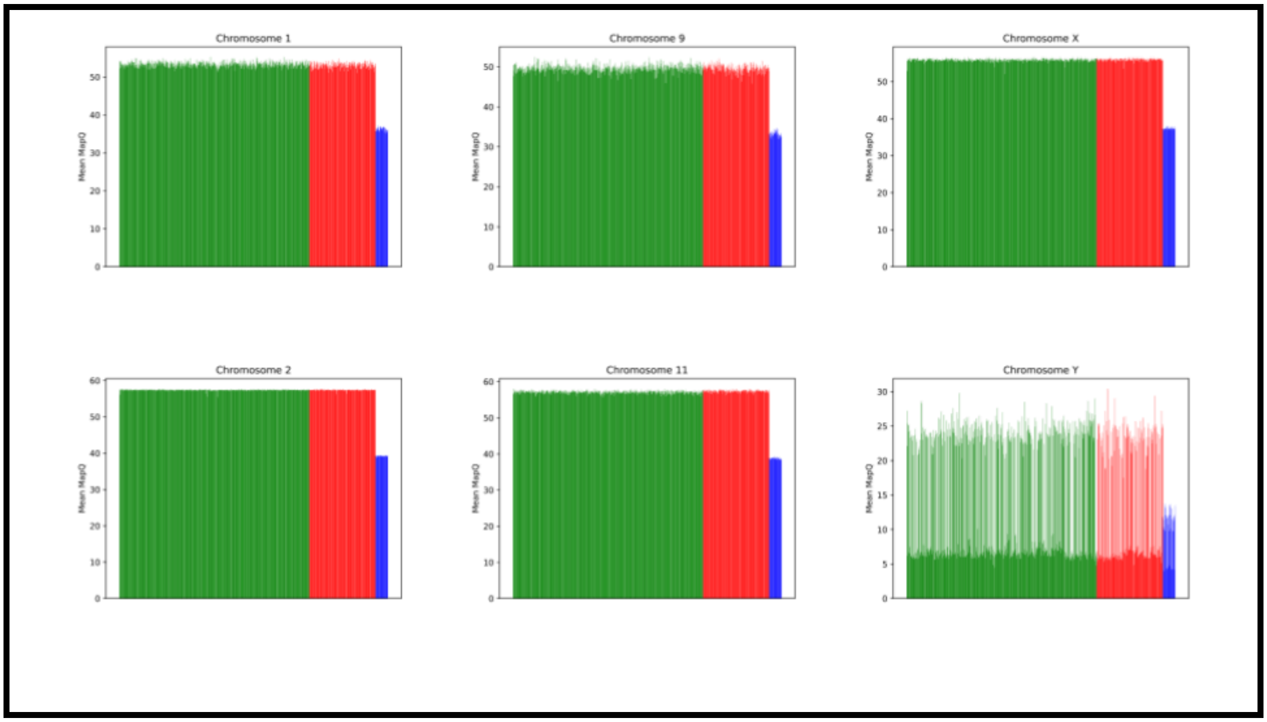


**Supplementary Figure 23: Average mapping quality against T2T-CHM13.** Mean mapQ of the read alignments to T2T-CHM13 across the 1KGP SAS (green), non-SAS (red) and SGDP SAS (blue sets) for chromosomes 1, 2, 9, 13, X and Y. Across all 24 autosomal and sex chromosomes, the SGDP individuals have consistently lower average mapQ.

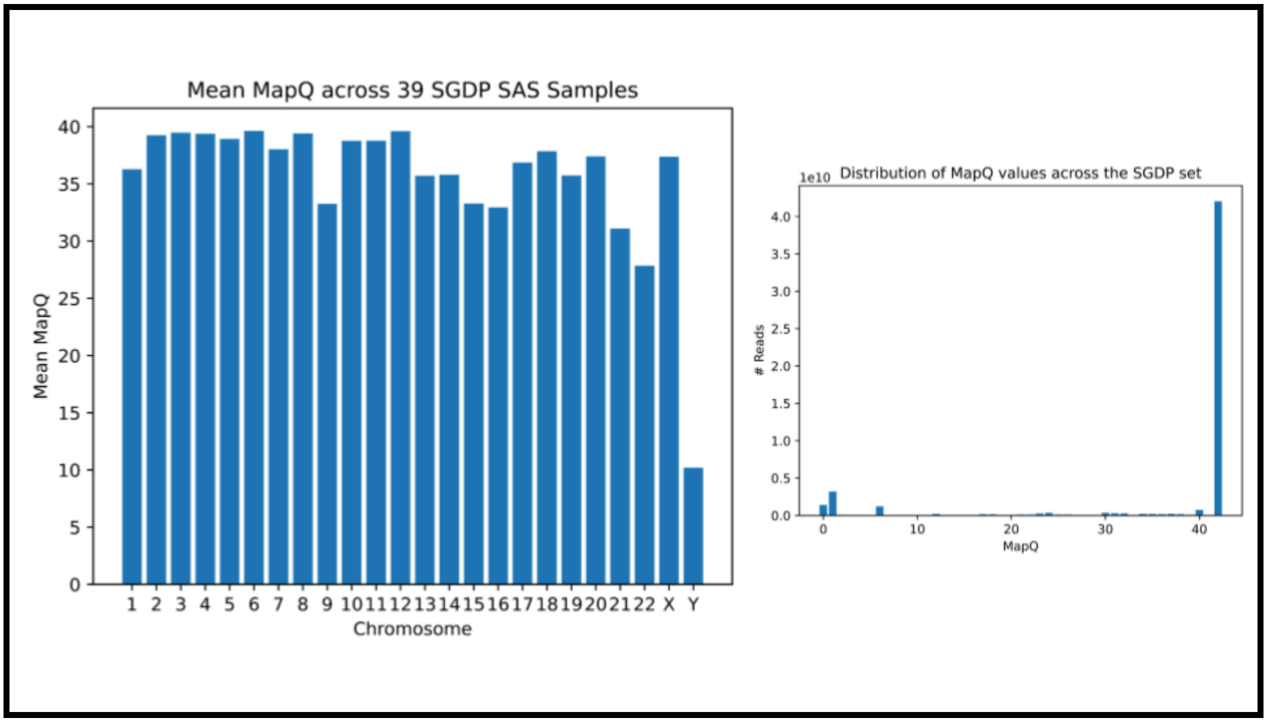


**Supplementary Figure 24: Mapping quality across SGDP chromosomes.** Mean MapQ across the autosomal and sex chromosomes in the SGDP set (**left**) and distribution of MapQ scores per read across the SGDP set (**right**). The majority of reads that do align align well, with a significant drop in mapping quality against chromosome Y.

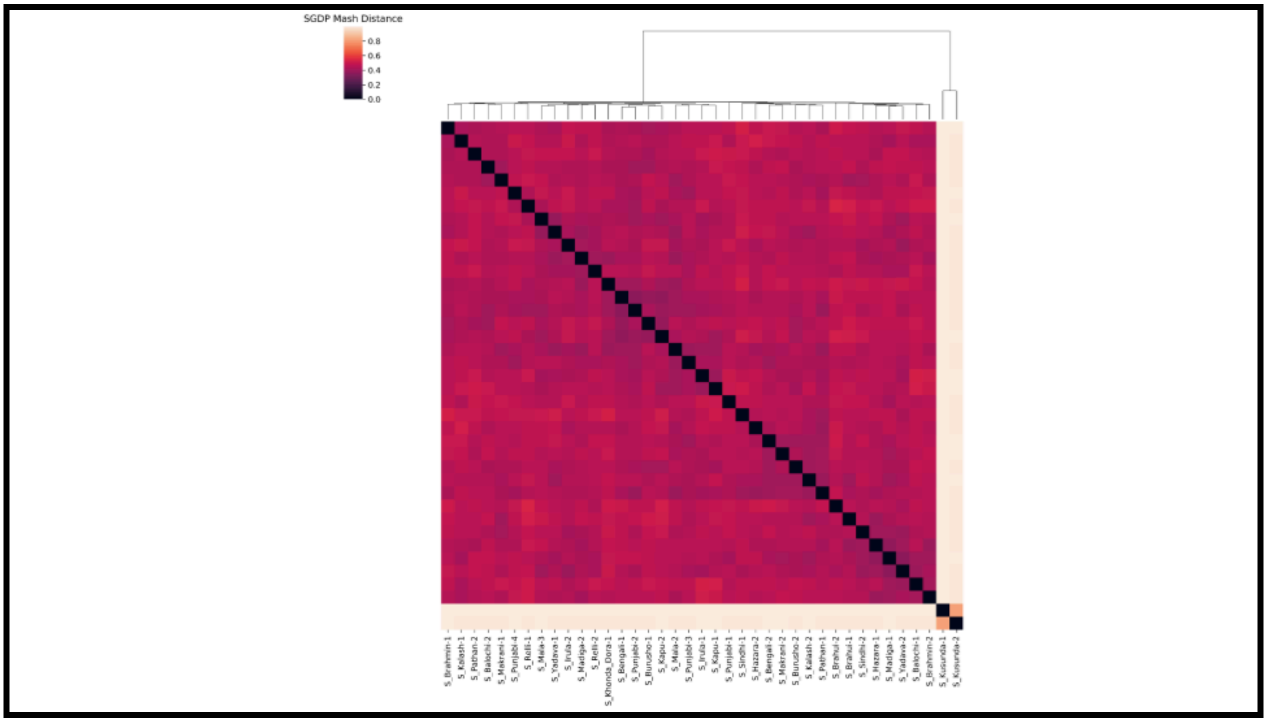


**Supplementary Figure 25: MASH-based clustering of SGDP contigs.** Mash distance-based clustering of the assembled unaligned read contigs across the 39 SGDP SAS individuals. The two Kusunda individuals are highly similar to each other, and significantly diverged from the rest of the set.

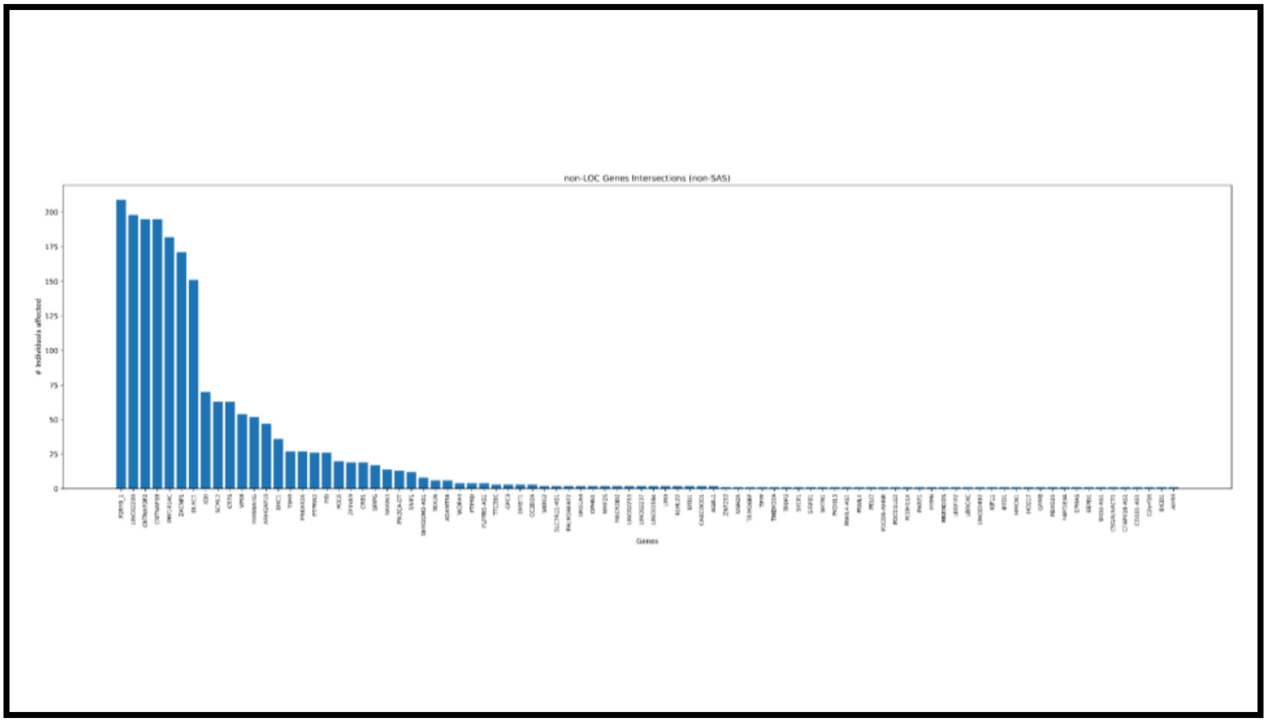


**Supplementary Figure 26: Non-SAS gene intersection distribution.** Number of non-SAS individuals (out of 210) that each of the 88 non-LOC genes intersected in the non-SAS set are intersected in. Full details and counts are in **Supplementary Table 1**.

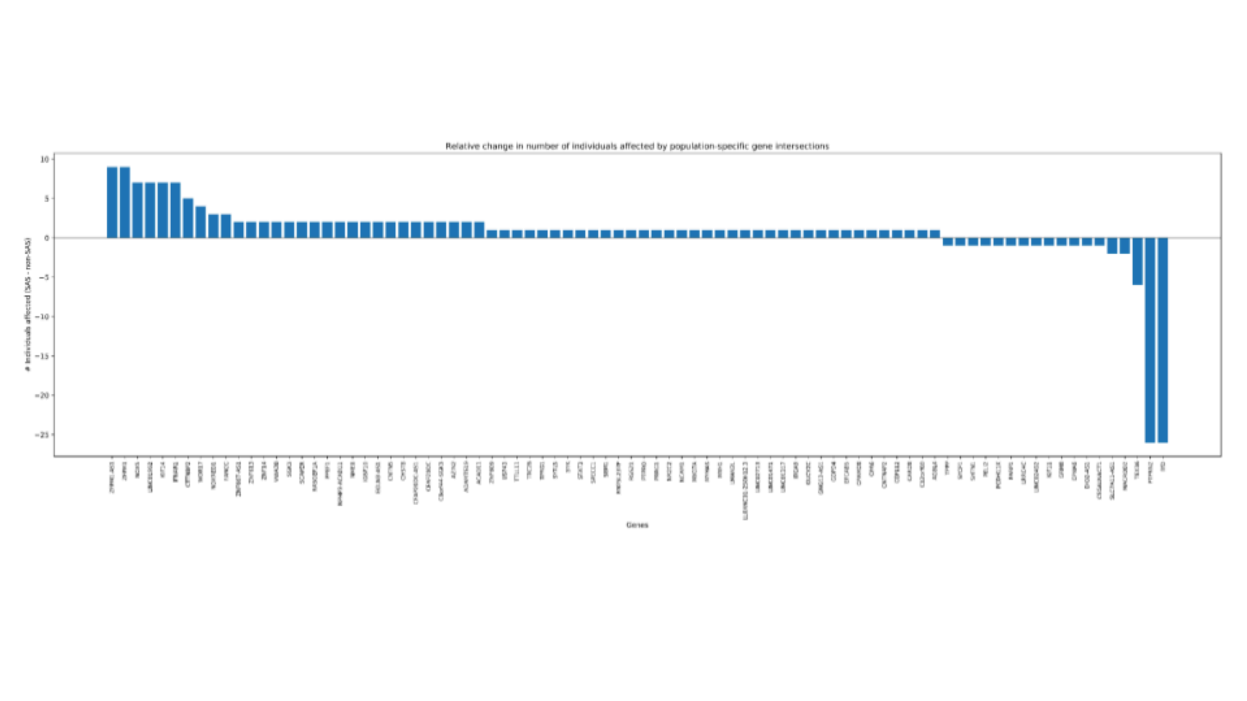


**Supplementary Figure 27: SAS vs non-SAS exclusive gene distribution.** Genes that only appear in the SAS set or non-SAS set, ordered based on the number of individuals in the corresponding set that they appear in. Does not include genes intersected by contigs from individuals in both sets. Full details and counts are in **Supplementary Table 1**.

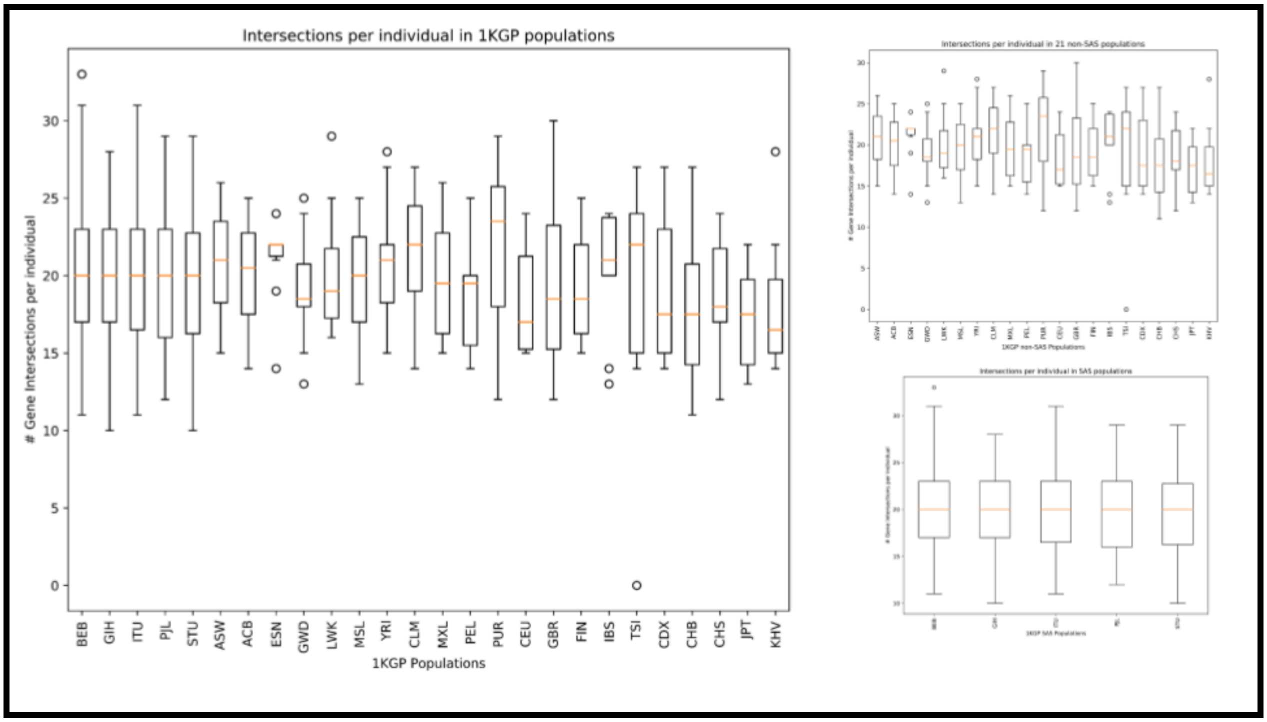


**Supplementary Figure 28: Gene intersections per individual across 1KGP populations.** Comparison of the number of gene intersections per individual **(a)** across the 26 1KGP populations, **(b)** in the 21 non-SAS populations, and **(c)** in the 5 SAS populations.

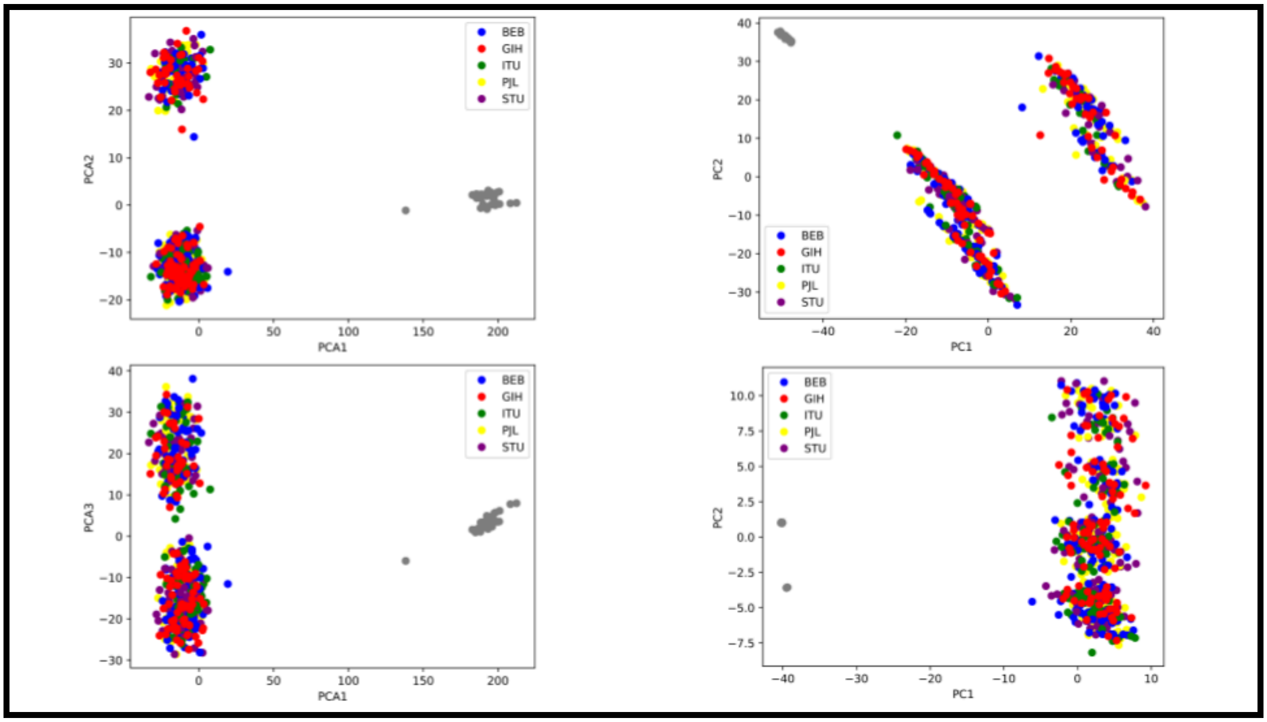


**Supplementary Figure 29: Clustering of SAS unaligned read contigs using PCA.** PCA plots using the presence/absence of 13,875 shared contigs in each of the 640 SAS individuals, across the first three PCs (**top and bottom left**). Adding constraints of only including contigs shared in 5 or more individuals and placed against T2T-CHM13 (**top right**) or further requiring the contigs to be present in both the 1KGP and SGDP subsets (**bottom right**) still largely recreates the same clusters. In both cases, the SGDP individuals cluster together, while the 1KGP SAS individuals split in roughly a 5:3 ratio.

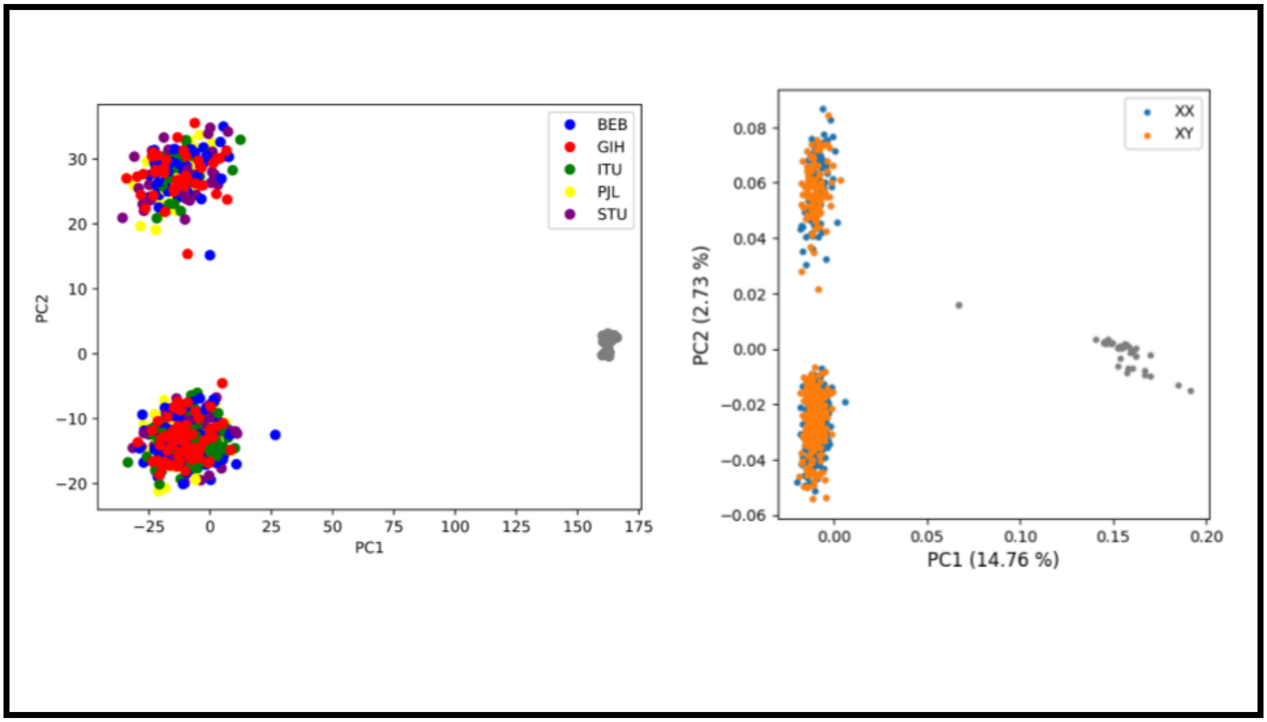


**Supplementary Figure 30: Further PCA Clustering Variations. Left:** PCA plots using the presence/absence of shared contigs that are placed against GRCh38 in each of the 640 SAS individuals, across the first two PCs. As with the PCA plots of placements against T2T-CHM13, the SGDP individuals cluster together, while the 1KGP SAS individuals split in roughly a 5:3 ratio. **Right:** PCA plots using the presence/absence of 13,875 shared contigs in each of the 640 SAS individuals, across the first two PCs, colored by karyotype. As seen in the plot, karyotype is not the driving force behind the split.

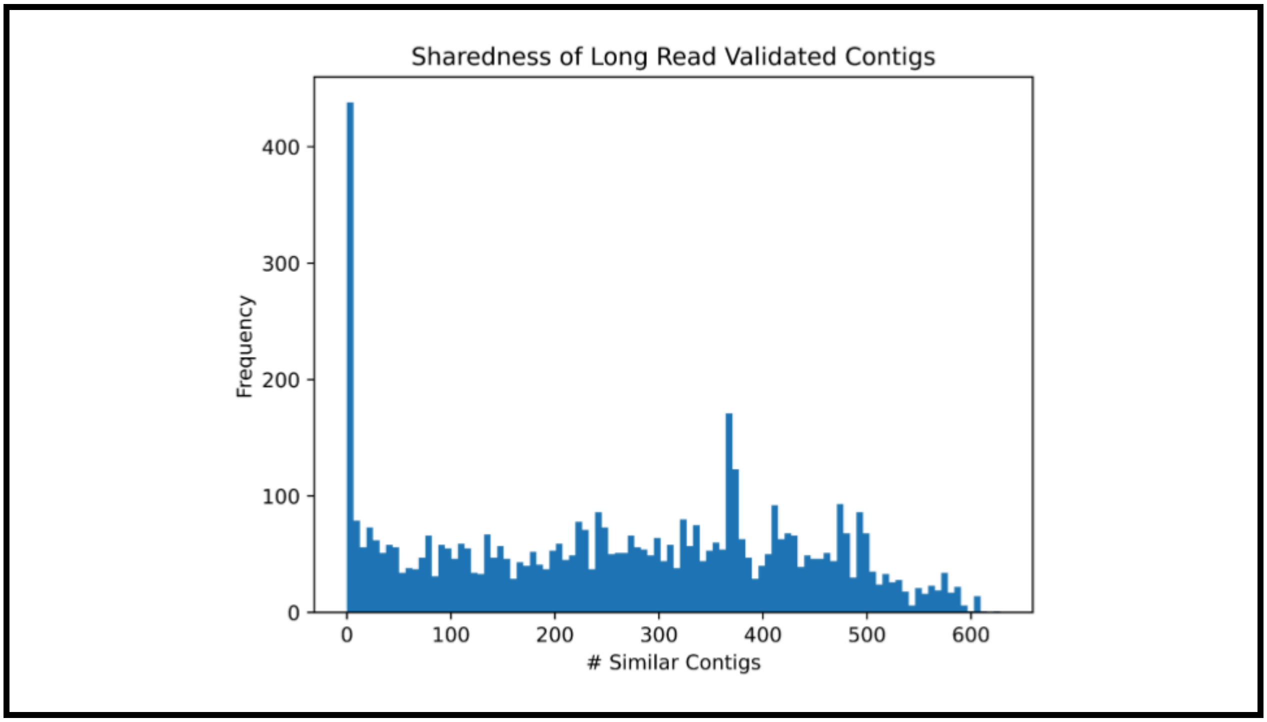


**Supplementary Figure 31: Sharedness of long-read validated contigs.** Sharedness of all long read-validated contigs in the 21 SAS individuals with long read data.

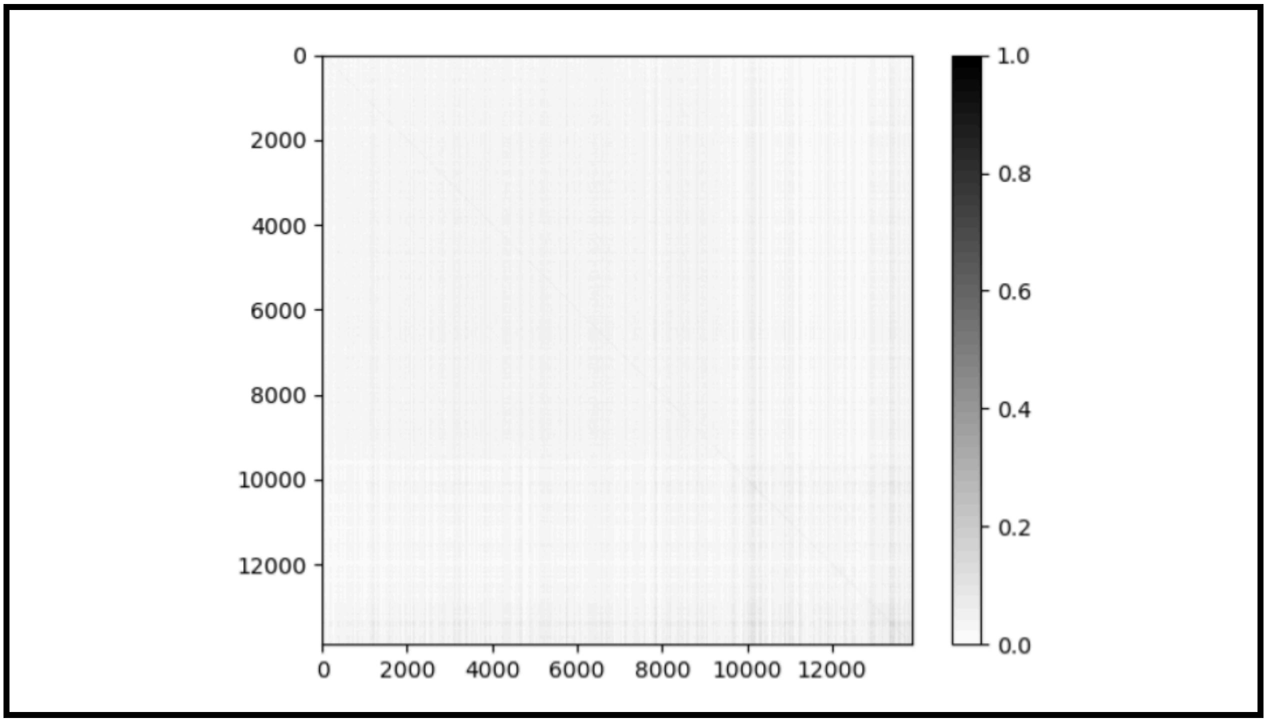

**Supplementary Figure 32: Possible associations across shared SAS contigs.** $R^2$ scores between the presence/absence of each of the 13,875 contigs across the 640 SAS individuals.

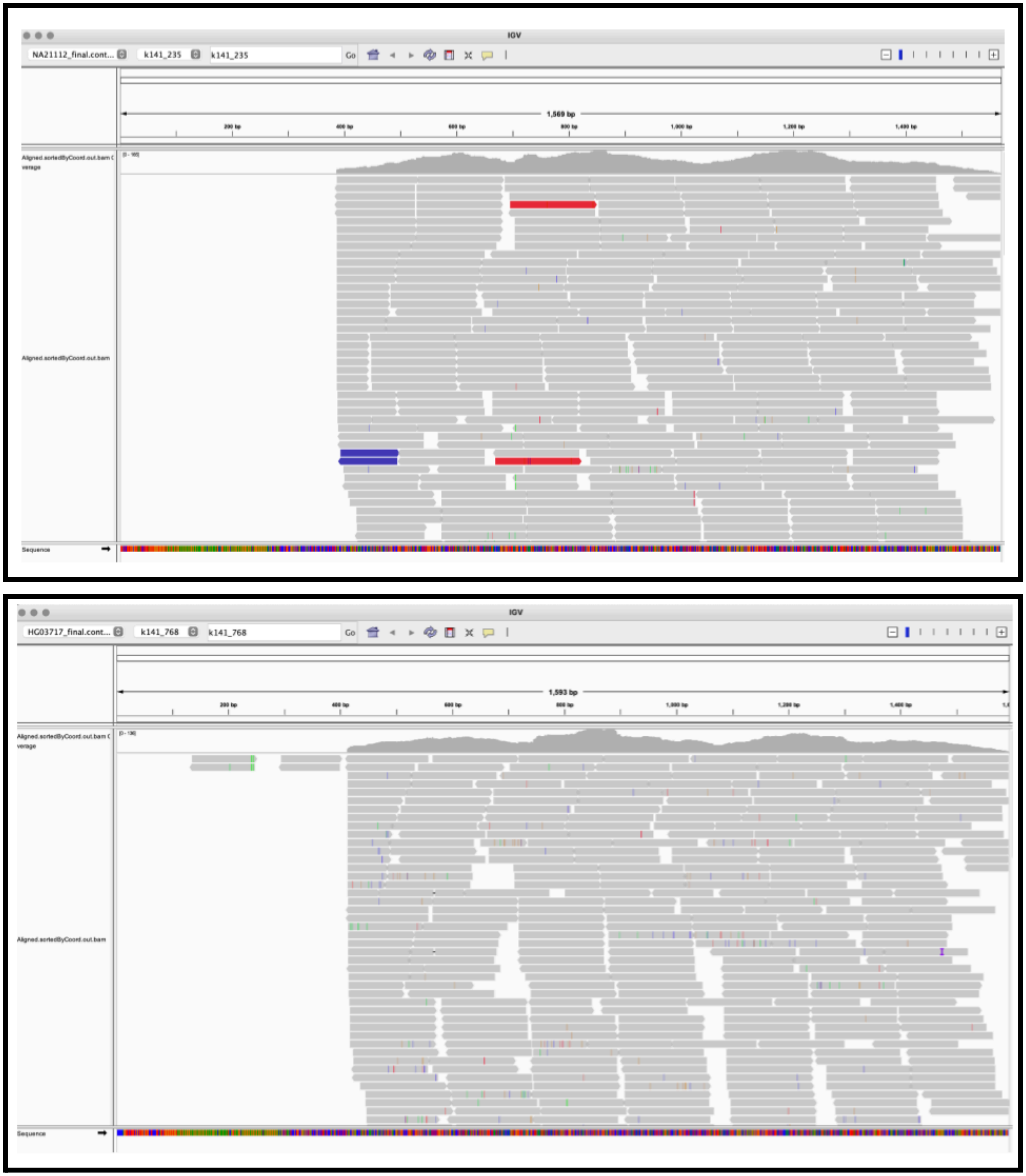


**Supplementary Figure 33: RNA-seq alignments to assembled contigs.** IGV screenshots of two contigs with possible exons, as evidenced by high number of RNA-seq reads aligned to regions of the overall contig. The two contigs are k141_235 in GM21112 **(top)** and k141_768 in HG03717 **(bottom)**, which are two of the clearest examples.

### Supplementary Notes

#### Supplementary Note 1: Alignment and Read Extraction

Alignment of input reads to chosen reference:

```
# Index reference
bowtie2-build ref.fa indexes/ref --threads ${THRDS}

# Align reads
bowtie2 -p ${THRDS} -x indexes/ref -1 ${RDS}_1.fastq -2
${RDS}_2.fastq > ${EXP}.sam

# Post-processing
samtools view -@ ${THRDS} -S -b ${EXP}.sam > ${EXP}.bam
samtools sort -@ ${THRDS} ${EXP}.bam -o ${EXP}.sorted.bam
```

Extraction of unaligned read pairs, and unaligned reads with a mapped mate:

```
# Get unaligned reads
samtools fastq -f 12 $ALGN -1 unmapped_R1.fq -2 unmapped_R2.fq -@
${THRDS}

# Get first reads that are unmapped but have a mapped mate
samtools fastq -f 68 -F 8 $ALGN > mateMapped_R1.fq -@ ${THRDS}

# Get second reads that are unmapped but have a mapped mate
samtools fastq -f 132 -F 8 $ALGN > mateMapped_R2.fq -@ ${THRDS}

# Get all unaligned reads into one file
echo "Extracting All Unaligned Reads"
samtools view -b -f 4 $ALGN > unmapped.bam -@ ${THRDS}
samtools sort unmapped.bam -o unmapped_sorted.bam -@ ${THRDS}
```

Extraction of reads based on quality value:

```
samtools view -h $ALGN -q 20 -o greater_than_q20.sam -U q20.sam -@
${THRDS}
samtools sort q20.sam -o q20_sorted.sam -@ ${THRDS}
samtools fastq q20_sorted.sam -1 q20_R1.fq -2 q20_R2.fq -@ ${THRDS}
```

#### Supplementary Note 2: Assembly and Filtering

Assembly of extracted reads using MEGAHIT:

```
megahit --num-cpu-threads 48 --out-dir ./megahit -1 unmapped_R1.fq -2
unmapped_R2.fq -r mateMapped_R1.fq, mateMapped_R2.fq
```

Extracting only contigs >1 Kb:

```
# Turn multiline into single line per sequence
awk '/^>/ { print (NR==1 ? "" : RS) $0; next } { printf "%s", $0 }
END { printf RS }' $ASM > collapsed.fa
# Extract 1Kb contigs
awk -v RS='>[^\n]+\n' 'length() >= 1000 {printf "%s", prt $0} {prt =
RT}' collapsed.fa > contigs_1Kb.fa
```

Running Centrifuge on the assembled contigs:

```
centrifuge -x centrifuge-1.0.3-beta/DB/p_compressed+h+v --report-file
/centrifuge.report -k 1 --host-taxids 9606 -f contigs_1Kb.fa >
centrifuge.output
```

Running BLAST on the assembled contigs:

```
blastn -db ref_prok_rep_genomes -query contigs_1Kb.fa -out
blast_output.out
```

The output files from Centrifuge and BLAST are then parsed to remove non-human sequence.

#### Supplementary Note 3: Placement

Minimap2 alignment of contigs to the chosen reference:

```
# Alignment
minimap2 -a -t 16 ref.fa megahit_unaligned_1kb_tagged.fa >
megahit_unaligned_1kb_tagged_mapped.sam
minimap2 -c -t 16 ref.fa megahit_unaligned_1kb_tagged.fa >
megahit_unaligned_1kb_tagged_mapped.paf
```

In addition to this, we use the mate-pair linking approach taken in the African Pangenome effort [15].

#### Supplementary Note 4: Intersection with Annotated Elements

Bedtools intersection with annotated elements:

```
# Bam to Bed conversion
samtools view -Sb megahit_unaligned_1kb_tagged_mapped.sam | bedtools
bamtobed -i - | bedtools sort -i - >
megahit_unaligned_1kb_tagged_mapped.sorted.bed

# Intersect with annotations file - keep details of intersection
bedtools intersect -a megahit_unaligned_1kb_tagged_mapped.sorted.bed
-b chm13v2.0_RefSeq_Liftoff_v5.1.gff3.gz -wb >
megahit_unaligned_1kb_tagged_mapped.intersect.bed

# No details, simple .bed file
bedtools intersect -a megahit_unaligned_1kb_tagged_mapped.sorted.bed
-b chm13v2.0_RefSeq_Liftoff_v5.1.gff3.gz >
megahit_unaligned_1kb_tagged_mapped.intersect_simple.bed
```

Bedtools intersection with GWAS sites (for direct overlap):

```
# Intersect with annotations file - keep details of intersection
bedtools intersect -a megahit_unaligned_1kb_tagged_mapped.sorted.bed
-b chm13v2.0_GWASv1.0rsids_e100_r2022-03-08.vcf.gz -wb >
megahit_unaligned_1kb_tagged_mapped.intersect_gwas.bed

# No details, simple .bed file
bedtools intersect -a megahit_unaligned_1kb_tagged_mapped.sorted.bed
-b chm13v2.0_GWASv1.0rsids_e100_r2022-03-08.vcf.gz >
megahit_unaligned_1kb_tagged_mapped.intersect_gwas_simple.bed
```

#### Supplementary Note 5: RNA-Seq alignments

STAR alignment of MAGE data to assembled contigs:

```
# Index the target sequences (combined set of all SAS MEGAHIT
post-filtering assemblies)
STAR --runThreadN 32 --runMode genomeGenerate
--limitGenomeGenerateRAM 125000000000 --genomeSAindexNbases 13
--genomeDir RNASeq/indexes/megahit_unaligned_1kb_tagged/
--genomeFastaFiles
RNASeq/combined_seqs/megahit_unaligned_1kb_tagged.fa

# Align
STAR --runThreadN 32 --genomeDir
RNASeq/indexes/megahit_unaligned_1kb_tagged/ --readFilesIn
SA_R1.fastq.gz SA_R2.fastq.gz --outFileNamePrefix RNASeq/alignments/
--readFilesCommand gunzip -c
```

#### Supplementary Note 6: BLAST querying

BLAST querying of unplaced and placed contigs:

```
# BLAST querying against nt db
# Unplaced
blastn -db nt -query megahit_unaligned_1kb_tagged_unplaced.fa -out
megahit_unaligned_1kb_tagged_unplaced.out -num_threads 32
# Placed
blastn -db nt -query megahit_unaligned_1kb_tagged_placed.fa -out
megahit_unaligned_1kb_tagged_placed.out -num_threads 32

# Extract the top 50 hits for each sequence
grep 'Query=' megahit_unaligned_1kb_tagged_unplaced.out -A 50 >
megahit_unaligned_1kb_tagged_unplaced_summary50.out
grep 'Query=' megahit_unaligned_1kb_tagged_placed.out -A 50 >
megahit_unaligned_1kb_tagged_placed_summary50.out

# Post process to analyze these hits
```

#### Supplementary Note 7: Long Read Assembly and Validation

Long read assembly using Flye:

```
# Convert unaligned read .bam to .fasta
samtools -@ 42 fasta sample.ONT.unaligned.bam > sample_ONT.fasta

# Flye assembly
mkdir flye_ONT_raw
flye --nano-raw sample_ONT.fasta --out-dir ./flye_ONT_raw --threads
42 -g 3.1g --debug | tee flye.log
```

Alignment of unaligned reads to long read assembly:

```
# Index
bowtie2-build sample_ONT.fasta indexes/sample_ONT --threads 36

# Align
(bowtie2 -p 24 -x indexes/sample_ONT -1 unmapped_R1.fq -2
unmapped_R2.fq -S unmapped.sam) 2> unmapped_ONT.log
```

Alignment of assembled contigs to long read assembly:

```
# Alignment in sam/paf format
minimap2 -a -t 32 ONT_assembly.fasta sample_contigs_1kb.fa >
unaligned_1kb_contigs.sam
minimap2 -t 32 ONT_assembly.fasta sample_contigs_1kb.fa >
unaligned_1kb_contigs.paf

# Counting total contigs
grep '>' sample_contigs_1kb.fa | wc -l
# Counting alignments with alignment length > 1Kb
awk '$11>1000' unaligned_1kb_contigs.paf | cut -f 1 | sort | uniq |
wc -l
```

#### Supplementary Note 8: All vs all alignment to identify shared sequences

```
# Self alignment with no limit on the number of secondary alignments
# Option 1
minimap2 -c -x asm5 -N1000 --cs -t 24
all_sas_megahit_unaligned_1kb_tagged_no_outliers.fa
all_sas_megahit_unaligned_1kb_tagged_no_outliers.fa >
all_sas_1kb_tagged_no_outliers_nolimit.paf

# Option 2
minimap2 -DP -cx asm5 -t 24
all_sas_megahit_unaligned_1kb_tagged_no_outliers.fa
all_sas_megahit_unaligned_1kb_tagged_no_outliers.fa >
all_sas_1kb_tagged_no_outliers_ava.paf
```

#### Supplementary Note 9: Pangenome Processing

```
# Alignment using GraphAligner
GraphAligner -g hprc-v1.0-minigraph-chm13.gfa -f reads_1.fastq
reads_2.fastq -a sample_ga.gam -x vg -t 20
GraphAligner -g hprc-v1.0-minigraph-grch38.gfa -f reads_1.fastq
reads_2.fastq -a sample_ga.gam -x vg -t 20

# Conversion to sam/bam/cram
vg surject -x hprc-v1.0-minigraph-chm13.xg -b sample_ga.gam >
sample_ga.bam
vg surject -x hprc-v1.0-minigraph-grch38.xg -b sample_ga.gam >
sample_ga.bam
```

Supplementary Note 10: Running Manta, Lumpy and PopIns2

```
## Manta ##

# prepare the analysis job
configManta.py \
  --bam sample~{bam_ext} \
  --referenceFasta ~{reference_fasta} \
  --runDir . &&

./runWorkflow.py \
  --mode local \
  --jobs ~{num_jobs} \
  --memGb $((~{num_jobs} * 2))

# inversion conversion, then compression and index
python2 /usr/local/bin/manta/libexec/convertInversion.py \
  /usr/local/bin/samtools \
  ~{reference_fasta} \
  results/variants/diploidSV.vcf.gz \
  | bcftools reheader -s <(echo "~{sample_id}") \
  > diploidSV.vcf

bgzip -c diploidSV.vcf > ~{sample_id}.manta.vcf.gz
tabix -p vcf ~{sample_id}.manta.vcf.gz



## Lumpy ##

# Cram to Bam conversion
samtools view -h -T ~{ref_fasta} ~{input_cram} | samtools view -b -o
${inputBam}.bam

# Get discordant reads
samtools view -h -@ ${threads} -F 1294 -u -b -h  ${inputBam} >
${inputBam}.temp && \
samtools sort -@ ${threads} -m 16G ${inputBam}.temp > discords.bam &&
rm ${inputBam}.temp

# Get split reads
samtools view -h -@ ${threads} ${bamToSplits} | \
/app/lumpy-sv/scripts/extractSplitReads_BwaMem -i stdin | \
samtools view -@ ${threads} -b -u - > ${bamToSplits}.temp && \
```

```
samtools sort -@ ${threads} -m 16G ${bamToSplits}.temp > splits.bam
&& rm ${bamToSplits}.temp

# Run LumpyExpress
lumpyexpress -B ${inputBam} -t ${threads} -S ${bamSplits} -D
${bamDiscords} -o ~{sample_name}_calls.vcf


## PopIns2 ##

# Link the reference genomes
ln -s CHM13.fa genome.fa
ln -s CHM13.fa.fai genome.fa.fai

# Assemble samples
popins2 assemble --sample sample1 sample1_CHM13.sorted.bam -t 24
...
popins2 assemble --sample sample10 sample10_CHM13.sorted.bam -t 24

# Merge
popins2 merge -r PopIns2/CHM13/ -di

# Contigmap
popins2 contigmap sample1 -t 24
...
popins2 contigmap sample10 -t 24

# Place
popins2 place-refalign
popins2 place-splitalign sample1
...
popins2 place-splitalign sample10
popins2 place-finish

# Genotype
popins2 genotype sample1
...
popins2 genotype sample10
```

#### Supplementary Note 11: Running RepeatMasker

```
# Default run
RepeatMasker --species Human
all_sas_megahit_unaligned_1kb_tagged_no_outliers.fa

# Run without masking low complexity DNA or repeats
RepeatMasker --species Human -nolow
all_sas_megahit_unaligned_1kb_tagged_no_outliers.fa

# Key output files are in
all_sas_megahit_unaligned_1kb_tagged_no_outliers.fa.masked,
all_sas_megahit_unaligned_1kb_tagged_no_outliers.fa.out and
all_sas_megahit_unaligned_1kb_tagged_no_outliers.fa.tbl
```
